# Temperature and genetic background drive mobilization of diverse transposable elements in a global human fungal pathogen

**DOI:** 10.1101/2025.05.19.654958

**Authors:** Anna I. Mackey, Vesper Fraunfelter, Samantha Shaltz, John McCormick, Callan Schroeder, John R. Perfect, Cedric Feschotte, Paul M. Magwene, Asiya Gusa

## Abstract

Transposable elements (TEs) are key agents of genome evolution across all domains of life. These mobile genetic elements can cause mutations through transposition or by promoting structural rearrangements. Stress conditions can amplify TE mobility, either by impairing TE suppression mechanisms or through stress-induced interactions between transcription factors and TE sequences, offering a route for rapid genetic change. As such, TEs represent an important source of adaptability within populations. To investigate the interplay between environmental stress and eukaryotic TE dynamics relevant to infectious disease, we examined how heat stress and host-mimicking medium (RPMI) affect TE mobility in the global human fungal pathogen *Cryptococcus neoformans*, using a collection of clinical and environmental isolates. Using a selection-based screen, we captured the mobilization of seven distinct mobile element families, encompassing diverse retrotransposons and DNA transposons, whose insertions conferred antifungal resistance. This includes a novel element, CNEST, which belongs to the CACTA, Mirage, Chapaev (CMC) supergroup. Heat stress at human body temperature (37°C) significantly increased the mobilization of a subset of these TEs, leading to higher rates of acquired antifungal resistance. Whole-genome assemblies revealed that, compared to retrotransposons, DNA transposons were hypomethylated and approximately uniformly distributed throughout the genome, features that may contribute to their frequent mobilization. We further assessed TE-driven genomic changes within hosts using serial isolates from patients with recurrent cryptococcal infections and from isolates passaged through mice. While we observed evidence of TE copy number changes near chromosome ends, we found no indication of TE-mediated alterations near gene-coding regions across any of the serial isolates. Finally, TE mobility was isolate- and strain-dependent, with significant variation even among clonally related strains collected from the same patient, emphasizing the role of genetic background in shaping TE activity. Together, these findings reveal a complex and dynamic relationship between environmental stress, genetic background, TE type-specific epigenetic regulation, and TE mobility, with important implications for adaptation and acquired antifungal resistance in *C. neoformans*.

**AUTHOR SUMMARY:** Transposable elements (TEs) are mobile segments of DNA that can “jump” from one place to another in a genome, causing genetic changes that can impact fitness and evolution. In this study, we investigate how different environmental conditions influence TE mobility in the pathogenic yeast *Cryptococcus neoformans*, which is globally distributed in the environment and causes life-threatening infections. We compared genetically diverse strains and found that growth at higher temperatures, specifically human body temperature (37°C), increases the mobility of some TEs. In contrast, other TEs were more active at lower temperatures (30°C), indicating distinct pathways of temperature regulation for different TE groups. By examining *C. neoformans* isolates from patients with recurrent infections, we observed genetic changes by one TE group that likely occurred during infection. We also found that TE groups are distributed differently across the genome and show varying levels of DNA methylation, a modification that can suppress their activity and control TE mobility in the cell. Overall, this work shows that environmental stresses, such as increased temperature, can accelerate genetic change by promoting TE mobilization. These changes may contribute to how *C. neoformans* adapts during infection or in the environment.

## INTRODUCTION

Transposable elements (TEs) are powerful drivers of genomic change and provide a mechanism for adaptive evolution in fungal pathogens [1–9]. TEs are mobile genetic elements that fall into two broad categories: retrotransposons, which mobilize via an RNA intermediate through a “copy-and-paste” mechanism, and DNA transposons, which mobilize via a DNA intermediate, typically through a “cut-and-paste” mechanism [10]. TE activity can lead to genomic instability through unchecked transposition or by providing homologous sequences throughout the genome that promote rearrangements [11]. However, TEs can also promote beneficial genetic changes, especially under changing or stressful environmental conditions, by modulating the expression of nearby genes, disrupting endogenous gene functions, or providing beneficial cargo [11–15]. Notably, various stressors can increase TE expression and mobilization rates in fungi [1,5,15–18]. TE-mediated genetic changes have been linked to multiple adaptive phenotypes, including increased fitness under heavy metal stress in *Schizosaccharomyces pombe* and nitrogen-limited conditions in *Saccharomyces cerevisiae* [1–5]. However, despite the demonstrated influence of TEs on adaptative genomic change, spontaneous transposition events are rarely observed in *S. pombe* and *S. cerevisiae* under many environmental conditions [19–22]. Therefore, the extent to which TEs contribute to the rapid adaptation of yeast to environmental stressors remains an open question.

Diverse TEs are present in *Cryptococcus neoformans*, a pathogenic yeast which primarily causes disease in immunocompromised individuals such as cancer patients, organ transplant recipients, and people living with HIV/AIDS [23–25]. *C. neoformans* is responsible for approximately 147,000 deaths annually and accounts for up to 20% of AIDS-related mortality [26,27]. It is a ubiquitous environmental yeast, and inhalation of spores or desiccated yeast cells can lead to pulmonary infection that may disseminate across the blood-brain barrier, causing cryptococcal meningitis [23]. Approximately 10 – 20% of patients diagnosed with cryptococcal meningitis experience recurrent disease, which can persist for years despite antifungal treatment [28]. In-host microevolution can result in adaptive genetic changes that enhance antifungal resistance, alter virulence traits, or increase overall pathogenicity, contributing to persistent infections and treatment failure [29–32]. Although in-host microevolution influences disease outcomes, the genetic basis of altered virulence traits and antifungal resistance often remains unknown, potentially because short-read sequencing and variant calling, the primary detection methods, are less suited for identifying structural rearrangements and TE polymorphisms [33].

During infection, *C. neoformans* encounters a variety of stressors during the environment-to-host transition. These include shifts in nutrient availability, pH, exposure to reactive oxygen species, and elevated temperature (37°C) [34]. Previous studies in the less pathogenic sister species *Cryptococcus deneoformans* have shown that heat stress significantly increases TE mobilization rates, exceeding those of single nucleotide polymorphisms (SNPs) and insertions/deletions (indels) by more than threefold per generation during in vitro passaging at 37°C [35,36]. These findings suggest that in stressful environments, TEs can be the primary drivers of genomic change. Indeed, TE mobilization in *C. deneoformans* has been observed during murine host infection and, in both *C. neoformans* and *C. deneoformans*, has been linked to the development of resistance to 5-fluorocytosine (5-FC), a first-line antifungal treatment [35–37]. A prior study in *C. neoformans* assessed whether heat stress increases mutation rates in the reference strain H99 and two additional clinical isolates, finding no association between heat stress and elevated mutation rates [38]. However, given the small number of isolates examined, it remains unclear whether heat stress can increase TE mobilization rates in *C. neoformans* and how it may affect mutation rates across additional isolates and lineages. Furthermore, it is unknown whether TEs contribute to genetic changes accumulated during persistent human infection in *C. neoformans*, which causes approximately 95% of cryptococcal infections [39].

*C. neoformans* is known to harbor a variety of TEs, including DNA transposons as well as long terminal repeat (LTR) and non-LTR retrotransposons [24,25]. Due to significant diversity within TE sequences, structures, and mobilization mechanisms, TEs are further subdivided using a hierarchical classification system [40]. TE orders (e.g., LTR) are subdivided into superfamilies (e.g., Ty1/Copia) based on defining structural or sequence features. For instance, DNA transposons that mobilize via a cut-and-paste mechanism typically contain a conserved DDE or DDD catalytic domain within the transposase, which can be used for identification [41]. To date, only a subset of TEs in *C. neoformans* have been classified to the superfamily level, including DNA transposons belonging to the Kyakuja/Dileera/Zisupton (KDZ) and HARBINGER superfamilies, and LTR retrotransposons from the Ty1/Copia and Ty3/Gypsy superfamilies [37,42,43]. Refining TE classification can provide insights into the mechanisms by which stress influences TE mobilization and genome evolution.

Because uncontrolled TE activity can cause genome instability, hosts have evolved mechanisms to suppress their mobilization [44–47]. The best-characterized TE suppression system in *C. neoformans* is RNA interference (RNAi). Through this pathway, transcripts targeted for silencing are processed into small interfering RNAs, which are loaded into the RNA-induced silencing complex to target homologous transcripts for degradation [43,48–50]. RNAi has been shown to silence double-stranded RNAs, transcripts with sub-optimal splicing signals, repetitive transgenes, and TEs in *C. neoformans* [38,43,48,50]. Unlike in the yeast *S. pombe*, RNAi is strictly involved in post-transcriptional silencing in *C. neoformans* and does not lead to transcriptional silencing by facilitating the assembly or maintenance of heterochromatin [43].

Prior studies have shown that loss-of-function mutations in key RNAi genes can lead to a hypermutator phenotype, driven by rampant TE mobilization, but only in strains with high TE burden [37,38]. TE mobilization may also be shaped by epigenetic context. LTR and non-LTR elements are enriched in centromeric and subtelomeric regions [38,42], where they co-localize with epigenetic features of silent chromatin, such as 5-methylcytosine DNA methylation and repressive histone modifications, including H3K27me3 and H3K9me2 [38,42,51–53]. These marks are also enriched in centromeric and subtelomeric regions, and are rarely detected in gene-rich chromosomes arms [38,42,51–53]. Despite this association, the regulatory effects of DNA methylation and heterochromatin on TEs in *C. neoformans* remain poorly understood, particularly regarding their influence on the mobilization of active TE families.

In this study, we investigated how infection-associated stressors, including elevated temperature and exposure to a host-mimicking medium (RPMI), influence TE mobilization in *C. neoformans*. Using a reporter gene-based approach, we examined TE mobility in genetically diverse clinical and environmental isolates, including representatives from three of the four major lineages (VNI, VNII, and VNBII) [54]. We found diverse, active TEs in *C. neoformans*, including several whose identity or mobility were previously unknown. TE mobility was temperature-dependent for all elements assessed, with heat stress significantly increasing transposition rates for a subset of elements. De novo telomere-to-telomere genome assemblies were used to examine TE-driven genetic changes in serially collected isolates from patients with recurrent cryptococcal meningitis. Using these assemblies, we annotated TEs to assess the genomic distribution and DNA methylation status of various transposable element types. DNA transposons were approximately uniformly distributed throughout the genome and had significantly reduced levels of DNA methylation compared to retrotransposons. Lastly, we found that TE copy number and mobility were highly isolate- and strain-dependent, suggesting that genetic background is a major determinant of TE-mediated genomic impacts. Together, these results highlight the critical roles of environmental factors, particularly temperature, in driving stress-induced genomic changes and adaptive evolution via transposition in the human fungal pathogen *C. neoformans*.

## RESULTS

### Diverse transposable elements mobilize in a subset of *C. neoformans* clinical isolates

To investigate transposable element (TE) mobilization dynamics in *Cryptococcus*, we obtained a set of 33 clinical isolates from 18 South African patients with recurrent cryptococcal meningitis, collected as part of the GERMS-SA surveillance study (Table S1) [55]. Isolates were collected by plating cerebrospinal fluid on non-selective medium and selecting a single colony at the time of diagnosis (incident) and after subsequent disease relapse(s), which occurred 57 – 238 days later, depending on the patient (Fig 1A, Table S1). Using short read sequencing, a prior study confirmed that isolates collected from the same patient differed by few single nucleotide polymorphisms (SNPs) and insertions/deletions (indels), and that these isolates formed monophyletic clades in phylogenetic analyses, suggesting that disease relapses were clonal and not due to reinfection [29]. This set of clinical isolates is genetically diverse, including *Cryptococcus neoformans* isolates belonging to three of the four major clades, VNI, VNII, and VNBII, and *Cryptococcus gattii* isolates belonging to clades VGI and VGIV (Table S1) [54,56].

**Fig 1.**
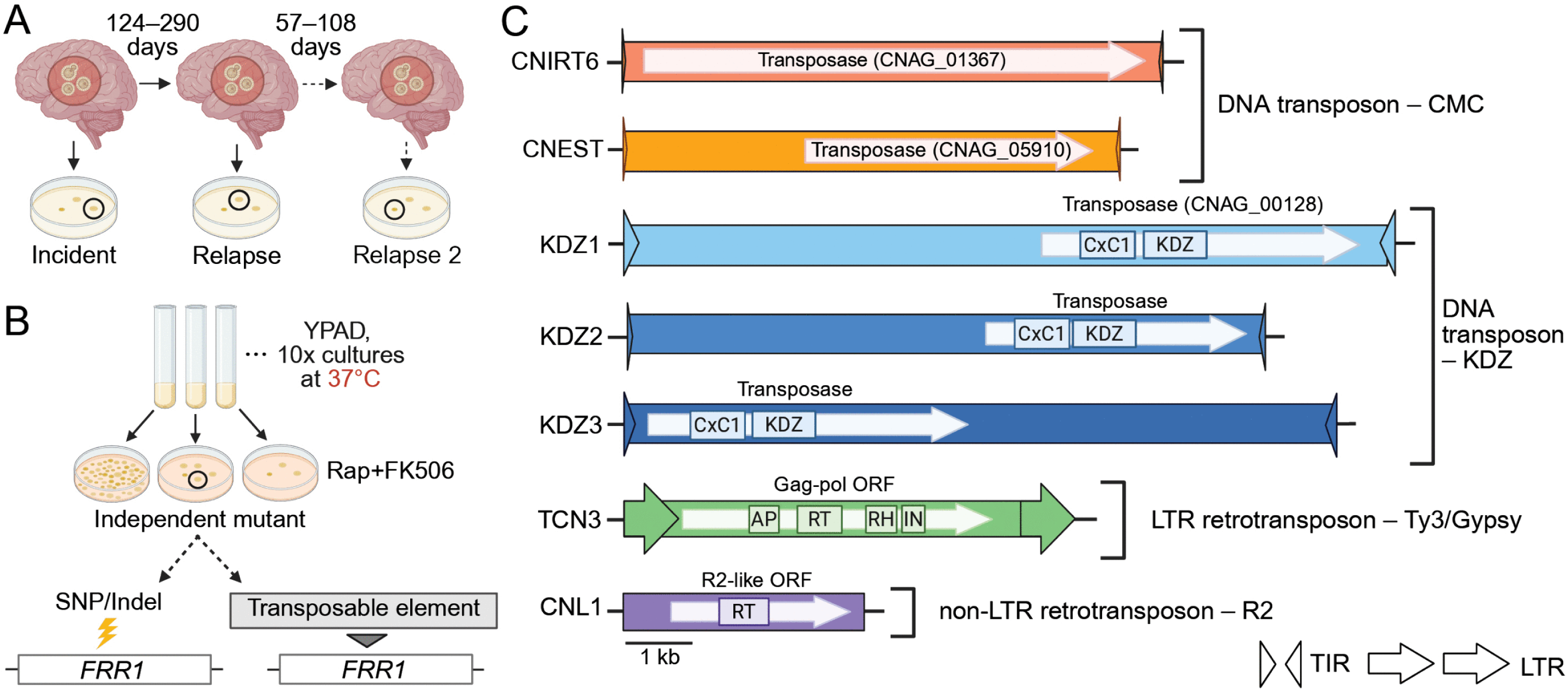
Diverse transposable elements (TEs) captured mobilizing in clinical isolates of *C. neoformans*. (**A**) Schematic of clinical isolate collection from patients with recurrent cryptococcal meningitis at disease onset and after subsequent relapse(s). Single colonies were isolated from patient cerebrospinal fluid on non-selective medium [55]. (**B**) Diagram of the preliminary screen using the “transposon trap” assay. Isolates were cultured in non-selective liquid YPAD medium at 37°C and plated on medium containing rapamycin and FK506 (rap+FK506) (N = 10 per isolate). Point mutations or TE insertions in the ∼934 bp *FRR1* (CNAG_03682) reporter gene confer resistance to rap+FK506. (**C**) Illustration of seven mobile TEs detected in *C. neoformans* isolates, including DNA transposons from the CMC (CACTA, Mirage, Chapaev) supergroup and KDZ (Kyakuja/Dileera/Zisupton) superfamily, and retrotransposons from the Ty3/gypsy and R2 superfamilies. Terminal inverted repeats (TIRs) and long terminal repeats (LTRs) are indicated by triangles and arrows, respectively. Genes within TEs that are predicted to facilitate mobilization are shown as white arrows. Functional domains are indicated, including KDZ (Kyakuja/Dileera/Zisupton transposase), CxC1 (predicted zinc chelating domain), AP (aspartic protease), RT (reverse transcriptase), RH (RNase H), and IN (integrase). TEs are drawn to scale. Created in BioRender. Gusa, A. (2025) https://BioRender.com/1psvbhx.

Prior work from our group showed that growth at 37°C increased TE mobilization rates in the less pathogenic species *Cryptococcus deneoformans* [35,36]. To investigate TE mobility under heat stress in *C. neoformans* and *C. gattii*, both recognized as priority fungal pathogens by the World Health Organization [57], we conducted a preliminary screen using a “transposon trap” assay that detects mutations in the ∼934 bp *FRR1* (CNAG_03682) reporter gene, a system previously used by our group and others [35,37,38]. Only mutations in *FRR1*, which encodes the shared drug target of the antifungals rapamycin and FK506 (rap+FK506), can confer resistance to both antifungals (Fig 1B) [58]. While mutations in other genes can confer resistance to rapamycin or FK506 individually, rap+FK506 resistance during a short growth period is most likely caused by a mutation in *FRR1*. This selection-based assay facilitates assessment of TE activity by requiring mutation analysis at only one locus. For the preliminary screen, incident and relapse isolates from the 18 South African cases, as well as the reference strain H99, were cultured in non-selective, nutrient-rich YPAD medium at 37°C and plated on medium containing rap+FK506 (Fig 1B). From a total of 37 isolates, ten independent rap+FK506 resistant mutants were collected per isolate, yielding 370 independent mutants. The *FRR1* gene was PCR amplified for each independent mutant to determine the causative mutation of drug resistance. Large DNA insertions, approximately 150 bp – 11 kb, detected by PCR amplification and indicative of TE insertions were confirmed by sequencing. Isolates from three of the 18 cases screened contained TE insertions disrupting the *FRR1* locus at 37°C: cases 7, 15, and 77, which belong to the VNBII, VNI, and VNII clades, respectively (Table 1). Isolates from these three cases were selected to assess the impact of temperature on drug resistance rates and TE mobility. TE mobilization was detected exclusively in *C. neoformans*; of the three *C. gattii* cases screened, only point mutations were identified in *FRR1*. Additionally, no TE insertions were identified in H99, consistent with prior studies [38]. Subsequent screening of rap+FK506-resistant mutants from cases 7 and 77 following growth at 30°C (see below) revealed additional mobile element families.

**Table 1.** Cases used for in-depth profiling of the impact of temperature on TE mobility, whole-genome assembly, and TE mapping.

| Case | Collection Point | Clade | Days After Incident Isolate Collection | TEs identified inserting in *FRR1* in YPAD |
| --- | --- | --- | --- | --- |
| 7 | Incident | VNBII | - | TCN3 |
| 7 | Relapse | VNBII | 137 | TCN3, CNL1, KDZ2, KDZ3 |
| 15 | Incident | VNI | - | CNIRT6 |
| 15 | Relapse | VNI | 166 | CNIRT6 |
| 15 | Relapse 2 | VNI | 223 | CNIRT6 |
| 77 | Incident | VNII | - | CNEST, KDZ1 |
| 77 | Relapse | VNII | 182 | CNEST, KDZ1 |
| 77 | Relapse 2 | VNII | 290 | CNEST |

In addition to the 370 independent rap+FK506 resistant mutants collected as part of the preliminary screen, we collected additional transposon trap mutants from cases 7, 15, and 77, along with an environmental isolate, for in-depth profiling of the effects of temperature (30°C and 37°C) and RPMI medium on TE dynamics. In total, we examined over 1,000 independent rap+FK506-resistant mutants and identified over 350 independent TE insertion events in *FRR1*. Elements were assigned to known families based on sequence similarity to *C. neoformans* TE family consensus sequences in the Repbase TE database, or to GenBank submissions when family-level consensus sequences were unavailable. Family membership was assigned based on the 80-80-80 rule: nucleotide similarity had to exceed 80% over at least 80 base pairs and cover at least 80% of the recovered sequence length [40]. An exception to this was the non-LTR retrotransposon CNL1, which shares only ∼60% nucleotide similarity with the CNL1 consensus sequence in Repbase. However, it exhibits high (> 99%) nucleotide similarity to an active mobile element previously classified as CNL1 in *C. neoformans* by Priest et al. (2022), so we have retained this name. In total, we identified seven diverse TE families mobilizing in *C. neoformans* clinical isolates (Fig 1C, Table 1). This includes an element belonging to a family that, to our knowledge, has not been identified in previous studies (CNEST), as well as elements not previously reported to be mobile (CNIRT6, TCN3, KDZ3) (Fig 1C, Table S2). These elements included DNA transposons, as well as long terminal repeat (LTR) and non-LTR retrotransposons.

### CNIRT6 transposition drives increased drug resistance rates at 37°C in case 15

To assess how heat stress impacts rap+FK506 drug resistance rates, fluctuation assays were performed on case 15 incident, relapse, and relapse 2 isolates. For all fluctuation assays, cultures were grown to saturation at 30°C or 37°C and the average cell population for each isolate and temperature condition was determined to ensure that differences in growth rates did not affect mutation rate estimates. All case 15 isolates displayed significant increases in rap+FK506 resistance rates during growth at 37°C compared to 30°C in YPAD medium, as indicated by non-overlapping 95% confidence intervals (21-, 12-, and 4-fold, respectively) (Fig 2A). The spectrum of mutations in *FRR1* was obtained by sequencing the *FRR1* locus of independent mutants, followed by mapping to the wild-type *FRR1* sequence for each isolate (Fig S1A). In cases where sequencing across the locus proved unsuccessful, junction PCR using TE-specific primers was used to determine the mutation type. Strikingly, many of the mutations in *FRR1*, up to 91% in the incident isolate at 37°C, were caused by insertions of a large (∼7.5 kb) DNA transposon with 99.7% nucleotide similarity to a putative element named CNIRT6 (GenBank: JQ277267.1) (Fig 2B). CNIRT6 was originally identified in *C. neoformans* strain JF289 through sequence analysis, however, this transposon has not been described in the literature and is absent from TE databases. Mapping of a subset of these insertions revealed CNIRT6 transposition into *FRR1* introns and exons, resulting in 3 bp terminal site duplications (TSDs) with variable sequences (Fig S2A, B). CNIRT6 insertion rates were significantly higher at 37°C compared to 30°C, increasing by 28-, 20-, and 16-fold in the incident, relapse, and relapse 2 isolates, respectively (Fig 2C). Notably, drug-resistance rates at 37°C and CNIRT6 mobility at both temperatures decreased progressively in relapse isolates collected 166 and 223 days after the incident isolate.

**Fig 2.**
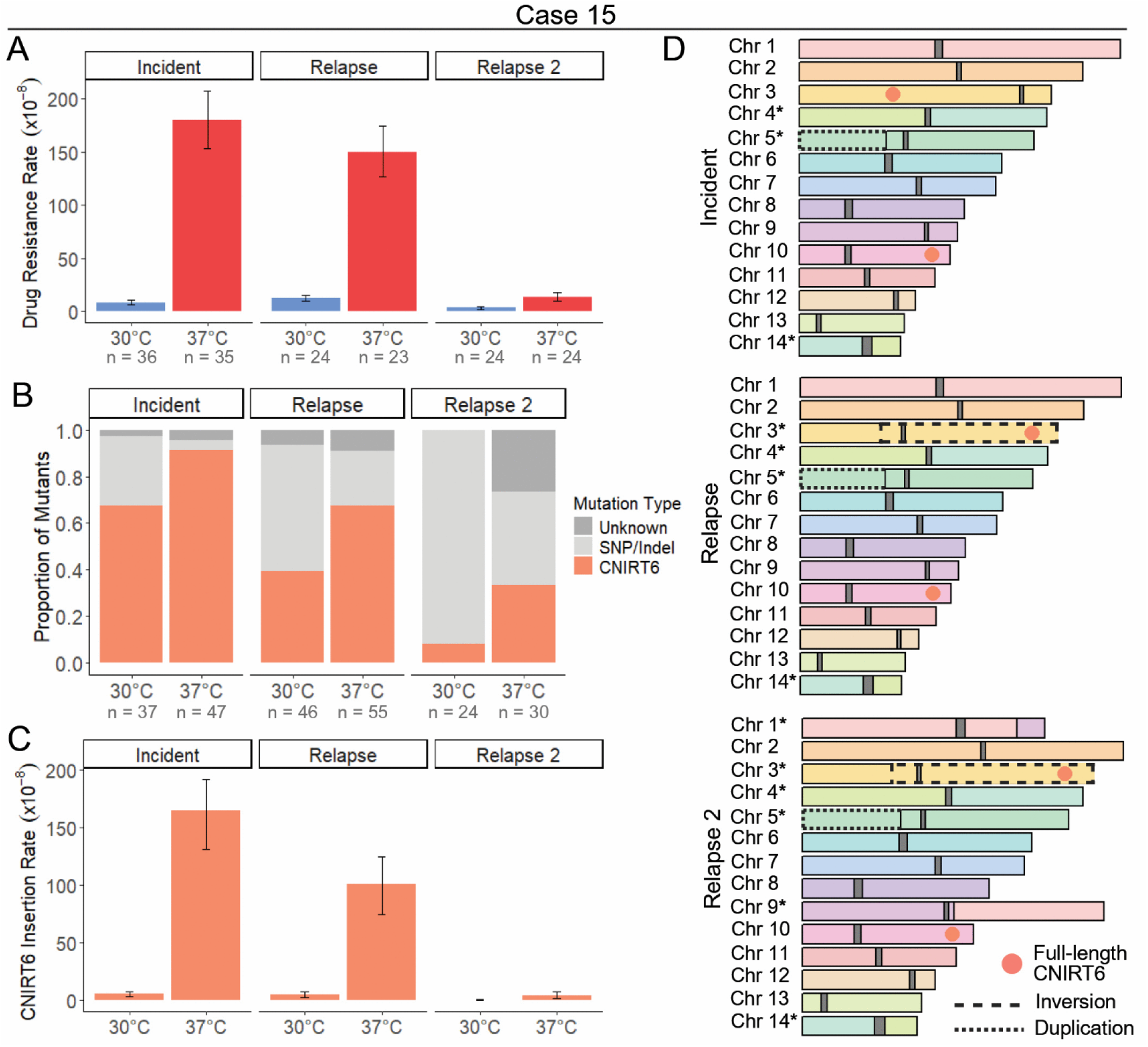
Mobilization of the DNA transposon CNIRT6 is induced by heat stress. (**A**) Rap+FK506 drug resistance rates per cell per generation following growth in non-selective YPAD medium at 30°C or 37°C (N = 23 – 36). (**B**) Mutation spectra in the *FRR1* gene for independent rap+FK506-resistant mutants. The mutation type “Unknown” indicates failure to amplify *FRR1* or absence of detectable mutations in its sequence (N = 24 – 55). (**C**) Insertion rate of CNIRT6 into *FRR1* per cell per generation. (**D**) Chromosome maps of the wild-type case 15 incident, relapse, and relapse 2 isolates. Full-length copies of CNIRT6 are shown as orange circles, and centromeres are shown as dark gray blocks. Chromosome synteny is indicated by chromosome color, with inversions and duplications outlined in dashed and dotted lines, respectively. All chromosomes with identified structural rearrangements are marked with an asterisk. Error bars in (**A**) and (**C**) represent 95% confidence intervals (CIs); rates are considered statistically different if error bars do not overlap.

Given the significant differences in CNIRT6 mobilization rates among case 15 isolates, we hypothesized that inter-isolate variability may be driven by differences in copy number, genomic position, or epigenetic regulation. To investigate these factors, and to assess all inter-isolate polymorphisms caused by mobile elements, we performed long-read sequencing to generate contiguous, telomere-to-telomere genome assemblies for the wild-type case 15 incident, relapse, and relapse 2 isolates, which were polished with Illumina reads (Table S3). 5-methylcytosine (5mC) DNA methylation was identified using Nanopore signal data. Given that all CNIRT6 insertions into *FRR1* appeared to be the same length by gel electrophoresis and shared high nucleotide similarity to one another (> 99%), the full-length CNIRT6 sequence, obtained from an insertion into *FRR1*, was used as the query in BLAST-like alignment tool (BLAT) [59] homology searches against case 15 genome assemblies to identify full-length copies (Fig S1A). For all BLAT homology searches predetermined cutoffs of nucleotide identity ≥ 90% and coverage ≥ 95% were used to call full-length copies. This analysis revealed two full-length CNIRT6 copies that are positionally conserved between incident and relapse isolates: one is an exact match to the CNIRT6 sequence identified inserting into *FRR1*, and the second differs by a single nucleotide substitution (Fig 2D). The nucleotide sequences of both CNIRT6 copies are identical between the incident, relapse, and relapse 2 isolates. No DNA methylation was detected in any full-length CNIRT6 copies across all case 15 isolates, as indicated by methylation frequencies < 0.75 at each CpG site, meaning fewer than 75% of long-read signal data supported a methylated call (Table S4). Therefore, differences in genomic copy number, chromosomal location, or DNA methylation do not appear to underlie differences in CNIRT6 mobilization rates between serial isolates.

To further characterize the active DNA transposon CNIRT6, we performed sequence analysis and found that this element was ∼7.5 kb in length and contained 24 bp terminal inverted repeats beginning with ‘CACTG’ (Fig 1C, Table S2). Though no CNIRT6 movement has been reported in the H99 reference strain, a homology search revealed two approximately full-length copies in H99 (identity ≥ 90% and coverage ≥ 95%), containing a ∼6.9 kb gene annotation (CNAG_01367). Repeat masking of the predicted protein sequence to assess similarity to known TEs using Repbase CENSOR revealed hits to transposases from TEs in the CACTA (also known as EnSpm) superfamily, which contain a ‘CACTA’ motif at their distal ends, resembling the ‘CACTG’ motif found in CNIRT6 [60]. To confirm that CNAG_01367 encodes a transposase and refine this classification, a multiple sequence alignment was performed using the amino acid sequences from CNAG_01367 and transposases from several cut-and-paste DNA transposons from the CMC (CACTA, Mirage, Chapaev) supergroup [41]. This alignment revealed a conserved DDE motif, which catalyzes the cut-and-paste mechanism of transposition, indicating that CNIRT6 represents a full-length element predicted to encode a transposase of the CMC supergroup (Fig S3A). To examine the relationship of CNIRT6 to other CMC transposons, an unrooted tree was constructed using transposase amino acid sequences representative of CMC families from animals, plants, and fungi. CNIRT6 formed a statistically supported clade (Bootstrap value 79%) with CACTA elements previously identified in other fungi, as well as plants and animals (Fig S3B). We conclude that CNIRT6 is a member of the CACTA superfamily within the CMC supergroup.

To comprehensively assess all genomic changes between case 15 isolates that may have been mediated by TEs other than CNIRT6, we performed synteny and read depth analyses, and used Synteny and Rearrangement Identifier (SyRI) [61], to define structural rearrangements and identify insertions, deletions, and duplications. We repeat masked insertions, deletions, and duplications, which revealed no evidence of transposition events between serial isolates, as indicated by the absence of sequence similarity to contiguous full-length TEs and transposition signatures such as TSDs. Relative to the genome reference strain H99, all three case 15 isolates contain a segmental duplication of ∼614 kb of the left arm of chromosome 5 and a translocation between chromosomes 4 and 14 where the breakpoint lies in the predicted centromeres (Fig 2D). When comparing the case 15 isolates to one another, we identified an inversion of ∼1.26 Mb on chromosome 3 present in both the relapse and relapse 2 isolates, and a translocation between chromosomes 1 and 9 present only in the relapse 2 isolate (Fig 2D).

Breakpoints for both inter-isolate rearrangements were outside of the predicted centromeres. No TE annotations were observed near rearrangement breakpoints, suggesting that another source of sequence homology or homology-independent DNA repair mechanisms mediated these rearrangements.

Lastly, given the robust activity of CNIRT6 in case 15 isolates grown in YPAD, we hypothesized that CNIRT6 may have mobilized during recurrent infections associated with other cases. To test this, we performed Southern blots on a subset of cases to identify putative transposition events between incident and relapse isolates. We generated a 551 bp CNIRT6-specific probe and utilized a restriction enzyme that cuts once within CNIRT6, outside of the probe site, and produces a range of fragment lengths separable by gel electrophoresis (Fig S4A). Consistent with whole-genome assembly comparisons, there was no difference in the hybridization band pattern for CNIRT6 among case 15 isolates (Fig S4B). However, several other patient cases displayed differences in CNIRT6 hybridization band patterns among serial isolates (Fig S4B). Case 1 was of particular interest, as it exhibited changes in the molecular weight of three bands, suggesting that these copies may occupy different genomic locations and indicating putative CNIRT6 transposition events between the incident and relapse isolates.

To investigate putative CNIRT6 transposition events in case 1 isolates, we generated whole-genome assemblies of the incident and relapse isolates and performed homology searches using full-length CNIRT6 as the query sequence. No full-length copies of CNIRT6 were detected in either genome; an identical number of truncated copies were present at the same genomic locations in both case 1 isolates (Fig S4C). Additionally, repeat masking of insertions, deletions, and duplications revealed no evidence of TE mobilization events between the isolates. To assess whether mutations that altered recognition sites of the restriction enzyme BspEI (which cleaves at the palindromic sequence ‘TCCGGA’) might explain differences in Southern blot banding patterns, we identified the closest BspEI sites upstream and downstream of CNIRT6 copies in the genomes. We found three instances where changes in BspEI sites led to the predicted production of CNIRT6-containing fragments of different lengths, which correspond to the observed band pattern differences, highlighting a limitation of this approach in detecting transposition events. Lastly, in the case 1 relapse isolate a translocation was detected between chromosomes 3 and 10, with the breakpoint located outside of the predicted centromere regions. There were no TEs near the breakpoint, suggesting that this rearrangement was not mediated by TE-derived sequence homology.

### CNEST transposition drives increased drug resistance rates at 37°C in case 77

For case 77, the relapse, and relapse 2 isolates displayed significant increases in rap+FK506 drug resistance rates during growth at 37°C vs 30°C (Fig 3A). Of mutations in *FRR1* observed after growth at 37°C, 29 – 82% were caused by insertion of a novel DNA transposon, which we named CNEST (*<u>C</u>ryptococcus <u>n</u>eoformans* <u>E</u>n<u>S</u>pm-like <u>T</u>ransposon; naming rationale described below) (Fig 3B). CNEST insertions appeared identical in length by gel electrophoresis and mapped to introns and exons in *FRR1* (Fig S1B, S5A). Insertions resulted in 2 bp ‘TA’ TSDs (Fig S5B). CNEST insertion rates were significantly higher for all isolates at 37°C; 22-, 83-, and 36-fold in the incident, relapse, and relapse 2, respectively (Fig 3C). The full-length CNEST sequence was ∼7.2 kb in length, with 33 bp TIRs beginning with ‘CACGG’ (Fig 1C, Table S2).

**Fig 3.**
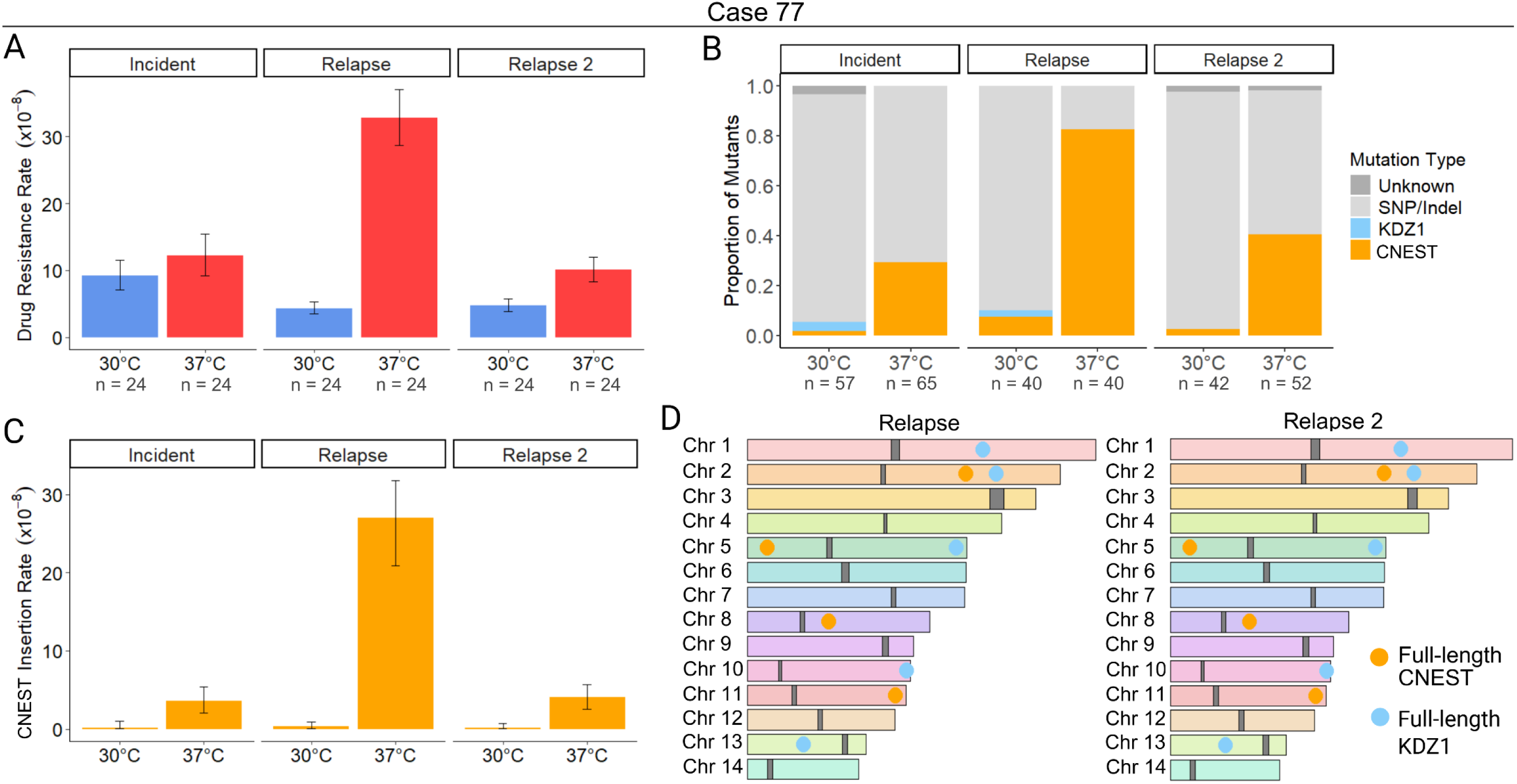
Mobilization of the DNA transposon CNEST is induced by heat stress. (**A**) Rap+FK506 drug resistance rates per cell per generation following growth in non-selective YPAD medium at 30°C or 37°C (N = 24). (**B**) Mutation spectra in the *FRR1* gene for independent rap+FK506-resistant mutants. The mutation type “Unknown” indicates failure to amplify *FRR1* or absence of detectable mutations in its sequence (N = 40 – 65). (**C**) Insertion rate of CNEST into *FRR1* per cell per generation. (**D**) Chromosome maps of the wild-type case 77 relapse and relapse 2 isolates. Full-length copies of CNEST (gold) and KDZ1 (light blue) are shown as circles, and centromeres are shown as dark gray blocks. Chromosome synteny is indicated by chromosome color. Error bars in (**A**) and (**C**) represent 95% confidence intervals (CIs); rates are considered statistically different if error bars do not overlap.

Interestingly, we also identified insertions of the DNA transposon KDZ1, recently characterized by Huang et al. 2024, which occurred at a lower frequency and only during growth at 30°C (Fig 3B) [37]. KDZ1 is an ∼11 kb element with 241 bp TIRs, and its insertion into *FRR1* resulted in 8 bp TSDs (Fig S5B, Table S2). The KDZ1 putative transposase contains a catalytic core affiliated with the Kyakuja/Dileera/Zisupton (KDZ) superfamily and a predicted Zn-chelating CxC1 domain, both of which have been previously described in fungi (Fig 1C) [62].

Assemblies of the case 77 relapse and relapse 2 isolates revealed conserved copy number and position of CNEST and KDZ1 by homology searches (identity ≥ 90% and coverage ≥ 95%) with 4 full-length copies of CNEST and 5 full-length copies of KDZ1 (Fig 3D). No DNA methylation was detected in any CNEST copies, as indicated by methylation frequencies < 0.75 at each CpG site from long-read signal data (Table S4). For KDZ1, a single copy contained one methylated CpG in the relapse (Table S4). In the relapse 2, two KDZ1 copies contained one methylated CpG. The additional methylated CpG in the relapse 2 isolate is ∼900 bp upstream of the predicted transposase start codon; however, a single methylated CpG seems unlikely to impact transcription of the transposase.

To further characterize CNEST, which has not previously been identified as mobilizing in the H99 reference strain, we performed a homology search, which revealed two approximately full-length copies in H99 (≥ 90% identity and ≥ 95% coverage), containing a ∼4.4 kb gene annotation (CNAG_05910). Repeat masking of CNEST and the predicted amino acid sequence of CNAG_05910 revealed no hits to TEs or transposases in the Repbase TE database.

However, given the identification of CNIRT6 as a member of the CMC supergroup, we predicted that CNEST may also encode a CMC transposase. Multiple sequence alignment of the CNAG_05910 amino acid sequence with transposases of the CMC supergroup revealed a well-conserved DDE catalytic core, confirming CNAG_05910 as a CMC-type transposase encoded within full-length CNEST (Fig S3A). Phylogenetic analysis places CNEST in a distinct clade from CNIRT6, forming part of a strongly supported clade (Bootstrap value 100%) containing unclassified transposases from diverse animals, including a transposase annotated as a CACTA (also known as EnSpm) element in the Pacific oyster (*Crassostrea virginica*) (Fig S3B). This clade is sister to clades of elements containing Chapaev and Mirage transposases from animals, forming a well-supported monophyletic group (Bootstrap value 86%) (Fig S3B). Thus, CNEST encodes a transposase defining a new superfamily within the CMC supergroup that is distinct but most closely related to Chapaev and Mirage superfamilies.

Lastly, no major genomic rearrangements were identified between the case 77 relapse and relapse 2 assemblies and no other TE-mediated genomic changes were detected by repeat masking, as all insertions, deletions, and duplications identified showed no significant sequence similarity to known TEs. Although mutation rate and spectra data are provided for the case 77 incident isolate, there was some variability in the growth of independent colonies and some inconsistency in mutation rate observed between independent fluctuation assays. The likelihood of genome instability in this isolate led to its exclusion from nanopore sequencing and genome assembly. In summary, two elements mobilized to mediate drug resistance in case 77: KDZ1, which was detected only at 30°C, and CNEST, which mobilized significantly more frequently at 37°C and is now characterized as a novel active TE in *C. neoformans* belonging to the CMC supergroup. Note that the KDZ superfamily, to which KDZ1 belongs, is completely distinct from the CMC supergroup, which includes CNEST and CNIRT6.

### Diverse TEs drive temperature and strain-specific TE mobilization dynamics in case 7

The case 7 incident isolate displayed a significantly higher rap+FK506 resistance rate following growth at 37°C compared to 30°C, while the relapse isolate showed no significant difference in drug resistance rates (Fig 4A). For the incident and relapse isolates at 37°C, ∼30% of mutations in *FRR1* were caused by insertions of the LTR retrotransposon TCN3, with both full-length and solo LTR insertions detected (Fig 4B, Fig S1C). TCN3 is ∼6.3 kb in length with 606 bp LTRs, and insertion into *FRR1* occurred in introns and exons, resulting in 5 bp TSDs of variable sequence (Fig 1C, Fig S6A, B, Table S2). TCN3 is a member of the Ty3/gypsy superfamily that has been previously characterized [63]. The TCN3 insertion we identified contains a ∼4.7 kb open reading frame with apparently intact Pol domains (reverse transcriptase, RNase H, protease, and integrase) (Fig 1C). Although we did not identify a Gag domain, the first 306 amino acids lack recognizable functional domains and correspond to the region that typically encodes Gag in Ty3/gypsy elements [63]. Moreover, the Gag gene is the most variable component of LTR retrotransposons, making it particularly difficult to detect [64]. These observations suggest that TCN3 likely represents an autonomous LTR retrotransposon (Fig 1C, Table S2). We recovered no TCN3 insertions at 30°C out of a total of 68 independent mutants screened, suggesting that the element may exclusively mobilize at higher temperatures (Fig 4B).

**Fig 4.**
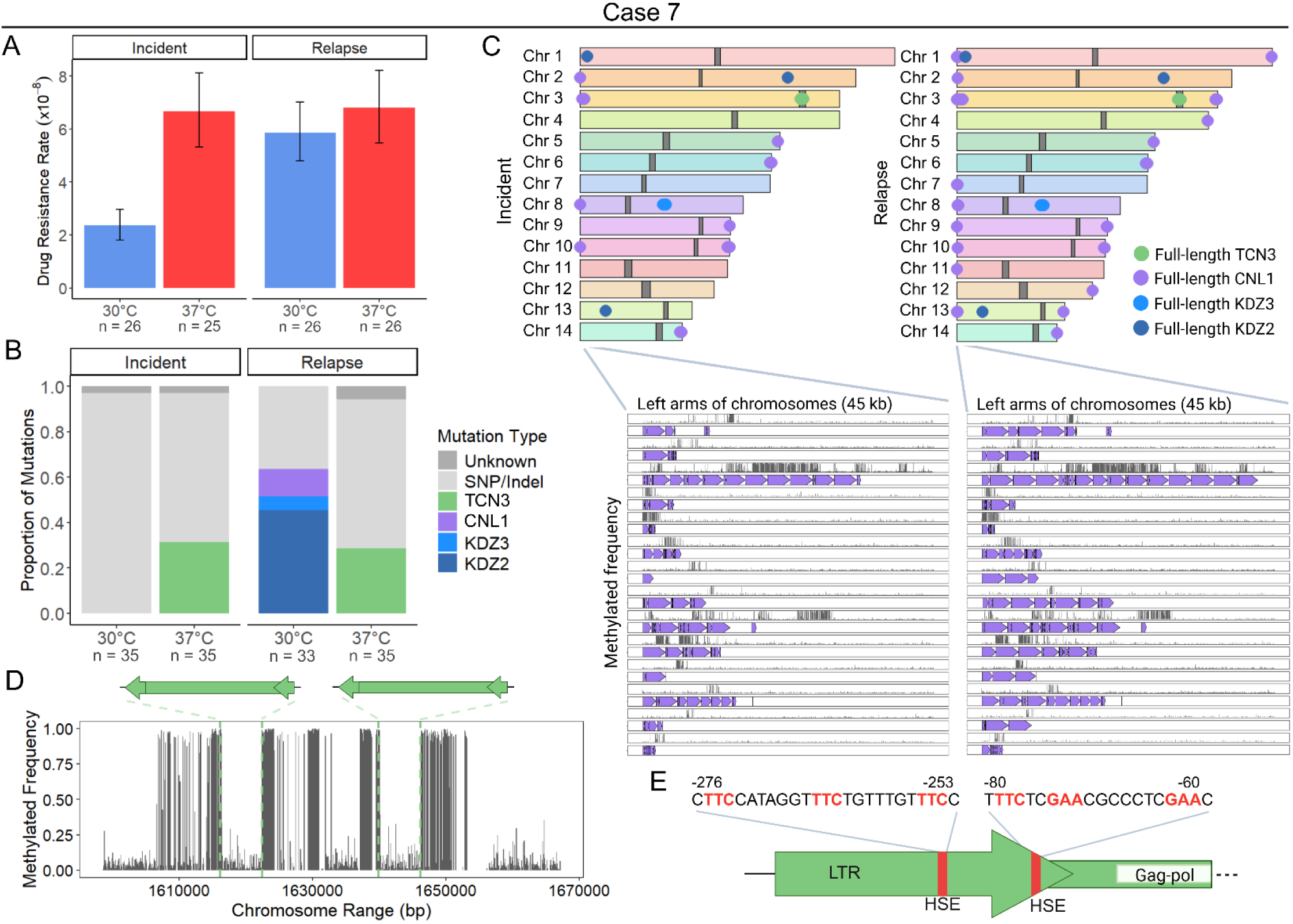
Temperature- and strain-specific TE dynamics in case 7. (**A**) Rap+FK506 drug resistance rates per cell per generation following growth in non-selective YPAD medium at 30°C or 37°C (N = 25 – 26). (**B**) Mutation spectra in the *FRR1* gene for independent rap+FK506-resistant mutants. The mutation type “Unknown” indicates failure to amplify *FRR1* or absence of detectable mutations in its sequence (N = 33 – 35). (**C**) Chromosome maps of the wild-type case 7 incident and relapse isolates. Full-length copies of TCN3 (green), CNL1 (purple), KDZ3 (blue), and KDZ2 (dark blue) are shown as circles, and centromeres are shown as dark gray blocks. Chromosome synteny is indicated by chromosome color. Insets show the 45 kb regions on the left arms of chromosomes in the incident and relapse isolates. Purple arrows indicate all CNL1 BLASTn hits; grey lines represent DNA methylation frequency from long reads, with the y-axis scaled from 0 to 1. (**D**) DNA methylation frequency plotted for the Chr 3 centromere region ±10 kb of the incident isolate. The position and orientation of full-length TCN3 copies is shown. (**E**) Diagram of HSE sequence motifs (red) identified in TCN3 LTRs. Position is shown relative to the gag-pol open reading frame (ORF) translation start site. Created in BioRender. Gusa, A. (2025) https://BioRender.com/ucajfxd.

Surprisingly, over 60% of *FRR1* mutations in the case 7 relapse isolate at 30°C were caused by insertions of diverse TEs, including the non-LTR retrotransposon CNL1 and the DNA transposons KDZ2 and KDZ3 (Fig 4B, Fig S1C). In contrast, mutations in the incident isolate at 30°C were exclusively SNPs/indels (Fig 4B). The longest CNL1 insertion we identified in *FRR1* was ∼3.5 kb in length and contained a ∼2.5 kb open reading frame with an apparently intact reverse transcriptase domain (Fig 1C, Fig S1C). This maximum insertion length matches previous reports of CNL1 transposition [38]. CNL1 insertions identified in *FRR1* ranged from 180 bp to ∼3.5 kb likely due to premature termination of reverse transcription which can occur during target primed reverse transcription (TPRT) [65]. CNL1 insertions occurred in the *FRR1* 5′ untranslated region (UTR) and in an exon, and were associated with TSDs of variable length (0 – 13 bp), consistent with findings from prior studies (Fig S6A, B) [38]. For KDZ2 (∼9.2 kb with 139 bp TIRs) and KDZ3 (∼10 kb with 169 bp TIRs), insertion into *FRR1* occurred in introns and exons and resulted in 4 – 8 bp TSDs (Fig 1C, Fig S6A, B, Table S2). The internal sequences of KDZ2 and KDZ3 encode putative transposases with KDZ and CxC1 domains (Fig 1C). Although TCN3 and KDZ3 were previously annotated, this study is the first to demonstrate their mobility.

Homology searches of case 7 genome assemblies using full-length TE sequences that inserted into *FRR1* as queries revealed conserved copy number and position of full-length TCN3 (2 copies), KDZ2 (2 copies), and KDZ3 (3 copies) between the incident and relapse isolates (Fig 4C). This indicated that the observed KDZ2 and KDZ3 activity in the relapse isolate was not due to an increase in copy number or change in genomic position. Additionally, no DNA methylation was detected on any full-length KDZ2 or KDZ3 copies in case 7 isolates, and no TE sequence polymorphisms were observed between serial isolates (Table S4). We did, however, identify an expansion of CNL1 copies in the relapse isolate, with the number increasing from 16 to 39 full-length copies in arrays in the subtelomeres (Fig 4C). CNL1 copies exhibited varying levels of DNA methylation (Fig 4C, Table S4). The expansion in CNL1 copy number may explain the observed CNL1 mobility in the relapse isolate, which was undetected in the incident isolate (Fig 4B). No genomic rearrangements or other TE-mediated changes, including alterations in copy number or position of TCN3 solo LTRs, were identified between the case 7 isolates.

Interestingly, we recovered 13 total CNL1 insertions in *FRR1*; however, all full-length CNL1 copies in case 7 genome assemblies are located at the ends of incident and relapse chromosome arms, flanked externally by telomeric repeats (“TAACCCCC,” or slight variations thereof), consistent with previous reports (Fig 4C) [24,66]. To further characterize CNL1, we performed a phylogenetic analysis of the reverse transcriptase domain from a full-length CNL1 insertion in *FRR1*, and the CNL1 consensus sequence from the Repbase TE database. This analysis revealed that CNL1 is more closely related to non-LTR elements of the R2 superfamily (Fig S3C), which are characterized by a single open reading frame and often insert into specific target sequences [67,68], in contrast to other non-LTR retrotransposons such as LINE1 elements which have weak target site specificity [69,70]. CNL1 was found to be most closely related to MoTeR, a sequence-specific telomeric-targeting element described in the fungal plant pathogen *Magnaporthe oryzae* (Bootstrap value 100%) [71]. Additionally, we observed similarity in the TSDs between independent CNL1 insertions in *FRR1*, suggesting potential sequence-specific target site preferences for this element (Fig S6A, B).

Centromeres in *C. neoformans* are typically enriched for LTR retrotransposons with > 95% of LTR retrotransposons, including TCN3, restricted to centromeric regions in the H99 genome reference strain, though the mechanism or selective pressure driving centromere targeting of insertions remains unclear [42]. Although endogenous TCN3 copies in case 7 isolates are restricted to centromeres, 21 independent TCN3 insertions were detected in *FRR1* (Fig 4B). Mapped insertions occurred at variable sequences and positions within *FRR1*, indicating a lack of strong sequence-specific insertion preference (Fig S6A, B).

*C. neoformans* centromeres are also enriched for DNA methylation, which is thought to contribute to repression of TEs present in these regions [51,52]. Therefore, we assessed whether the centromeric copies of TCN3 were methylated. If methylated, demethylation and subsequent derepression of TCN3 under heat stress in vitro might explain the observed TCN3 mobility at 37°C in case 7 isolates [72]. We found that the full-length copies of TCN3 in the case 7 isolate genomes reside in hypomethylated regions within the predicted centromere region of Chr 3 for the incident and relapse (data for the incident isolate shown) (Fig 4D). While five to seven 5mC DNA methylation marks were observed in full-length TCN3 copies, they are located at the end of the 3’ LTR (Fig 4D, Table S4). This region is not typically transcribed, making the observed DNA methylation unlikely to impact TCN3 transposition [73]. This suggests that loss of DNA methylation at full-length TCN3 copies is unlikely to explain the differential activity of TCN3 observed in vitro, where transposition occurred only at 37°C.

Our group previously identified the LTR-retrotransposon TCN12 in *Cryptococcus deneoformans* as heat stress-activated, mobilizing almost exclusively at 37°C [35,36]. In *C. neoformans*, heat shock binding elements (HSEs) in the promoter region of heat-responsive genes have been shown to be bound by the transcriptional activator heat shock factor 1 (Hsf1) during heat stress [74,75]. A putative HSE was previously identified in the TCN12 LTR [36], and upon scanning the TCN3 LTR, we identified two putative HSEs (Fig 4E). This suggests that the elevated mobilization rates of TCN3 and TCN12 at 37°C may result from Hsf1 binding and subsequent upregulation.

### TE mobility is altered in RPMI medium

To more closely mimic the environment *C. neoformans* experiences in-host and assess impacts on transposition, we compiled the spectra of mutations in the *FRR1* reporter gene for cases 7, 15, and 77 following growth in chemically defined RPMI tissue culture medium at 37°C [76]. We additionally tested the impact of RPMI medium at 30°C for case 7 isolates given the activity of CNL1, KDZ2, and KDZ3 observed in the relapse isolate at 30°C in YPAD medium (Fig 4B). RPMI medium was supplemented with less glucose (0.2%) compared to YPAD (2%) and had a defined starting pH of 7.0. For four of the six isolates, the same TEs were identified as mobile in both YPAD and RPMI medium (Fig S7A, B, C). In case 77 and case 7 incident isolates, CNL1 and KDZ3, respectively, were observed inserting into *FRR1*, events not previously detected in YPAD medium (Fig. S7B, S7C).

We next assessed differences in the frequency of TE insertion events between RPMI and YPAD medium. In the case 15 relapse isolate, there was a significant difference in TE insertions into *FRR1*, with fewer insertions occurring in RPMI compared to YPAD at 37°C (two-tailed Fisher’s Exact test, *p* = 5.62e-6) (Fig S7A, Fig 2B). Interestingly, the use of RPMI medium altered TE dynamics in an isolate- and temperature-specific manner in case 7 isolates (Fig S7C, Fig 4B). Differences between RPMI and YPAD medium include: (1) a significant decrease in TCN3 insertions in the relapse at 37°C in RPMI, with no insertions detected (two-tailed Fisher’s Exact test, *p* = 0.00905), (2) a significant increase in KDZ3 insertions in the incident at 30°C in RPMI, with no insertions occurring in YPAD (two-tailed Fisher’s Exact test, *p* = 2.56e-4), and (3) a significant increase in CNL1 insertions in the relapse at 30°C in RPMI (two-tailed Fisher’s Exact test, *p* = 0.0261), highlighting the complex relationship between environment, genetic background, and TE mobility in case 7 isolates. Across all cases assessed, growth in RPMI medium altered either the TE families observed mobilizing or the frequency of insertion events for at least one isolate.

### CNEST heat stress-activation is conserved in a geographically and temporally distinct environmental isolate

Homology searches revealed full-length CNEST and KDZ1 copies in a previously published *C. neoformans* VNII genome, the same lineage as the case 77 isolates, sequenced from an environmental sample collected from cockatoo droppings in Boston in 2000 (designated cockatoo guano isolate) [31,77]. While the cockatoo guano and case 77 isolates both belong to the VNII clade, they are geographically and temporally distinct, as the case 77 isolates were collected eight years later in South Africa. However, a pairwise whole-genome comparison revealed that the isolates are highly genetically similar, differing by 1,610 SNPs and indels across the genome, corresponding to approximately 0.0085% divergence at the nucleotide level. To better understand mobility of CNEST and KDZ1 in a different strain background, we assessed rap+FK506 drug resistance rates and the spectra of mutations in *FRR1*. Indeed, cockatoo guano isolate drug resistance rates were significantly elevated during growth at 37°C compared to 30°C (Fig 5A). Approximately 20% of mutations in *FRR1* at 37°C were CNEST insertions, while no CNEST insertions were observed at 30°C, strongly supporting that CNEST transposition is induced by heat stress in the cockatoo guano isolate (Fig 5B). To assess the genomic context of CNEST and KDZ1, we performed synteny analysis, which demonstrated that both the copy number and genomic position of CNEST and KDZ1 were conserved between the cockatoo guano and case 77 isolates (Fig 5C). This may suggest that mobilization of CNEST and KDZ1 in the VNII lineage is infrequent in situ, that mobilization events are typically detrimental to fitness, or that there is selective pressure to maintain both the copy number and genomic context of these elements, all possibilities that will require further investigation.

**Fig 5.**
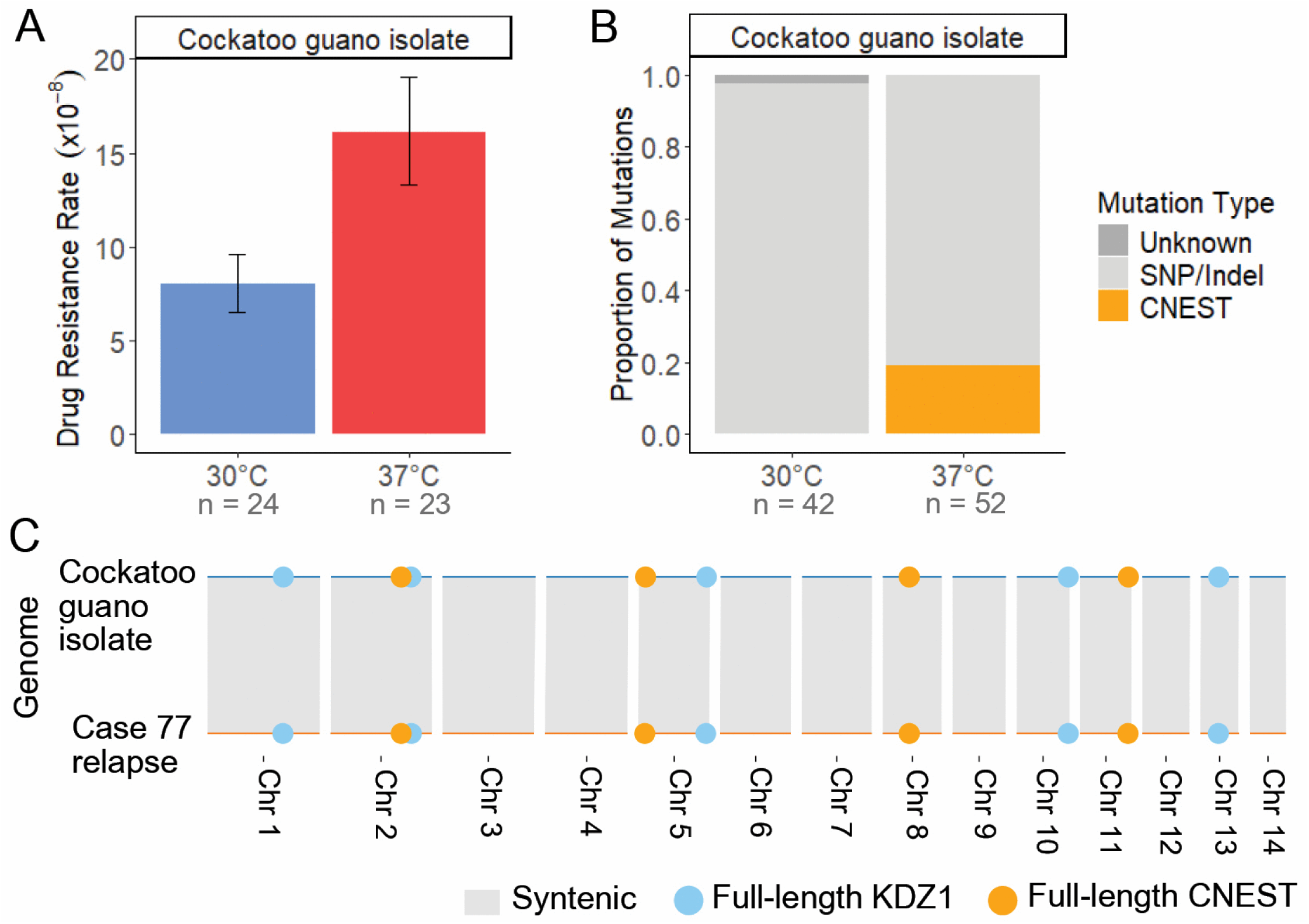
Heat stress-induced CNEST activation is conserved in an environmental isolate. (**A**) Rap+FK506 drug resistance rates per cell per generation following growth in non-selective YPAD medium at 30°C or 37°C (N = 23 – 24). Error bars represent 95% confidence intervals (CIs); rates are considered statistically different if error bars do not overlap. (**B**) Mutation spectra in the *FRR1* gene for independent rap+FK506-resistant mutants. The mutation type “Unknown” indicates failure to amplify *FRR1* or absence of detectable mutations in its sequence (N = 42 – 52). (**C**) Chromosome-level synteny between the cockatoo guano and case 77 relapse isolates. Full-length copies of CNEST (gold) and KDZ1 (light blue) are shown as circles.

Copy number analysis of other mobile TEs identified in this study revealed two full-length copies of KDZ3 in the case 77 genomes and one full-length copy in the cockatoo guano isolate, located at a conserved site shared with one of the case 77 copies (Fig S8A). The second KDZ3 copy in the case 77 relapse isolate, located on Chr 12, appears to result from a transposition event, as synteny flanking this copy is maintained between the two isolates and 8 bp TSDs were identified (Fig S8A, B). Repeat masking of insertions and deletions also revealed evidence of an additional TE family, Gypsy-1_CrGa, originally identified in *C. gattii* [78], whose copy number differs between the case 77 relapse and cockatoo guano isolates (Fig S8A). Two Gypsy-1_CrGa transposition events were detected in the cockatoo guano isolate and one in the case 77 relapse genome (Fig S8A). All insertions of this element occurred within predicted centromeres and generated 6 bp TSDs of variable sequence (Fig S8B). Other insertions and deletions, which commonly occurred in the centromeres, contained TE sequences but were likely not mediated by transposition, as they involved multiple or fragmented TE copies and lacked transposition signatures such as TSDs. Thus, while KDZ1 and CNEST copy number and position are conserved between these isolates, transposition events involving other TE families have occurred.

A recent study passaged the cockatoo guano isolate twice through A/J mice via retro-orbital intravenous injection, collecting a total of ten isolates from the brains and lungs after the second passage [31]. The cockatoo guano isolate was also the point-source isolate for a case of cryptococcal meningitis, from which a single isolate was collected from the patient (PU isolate) [77]. To assess whether transposition of the two highest copy number TEs located outside of the predicted centromeres, CNEST and KDZ1, occurred during murine passage or human infection, we performed Southern analysis using 551 bp CNEST- and 567 bp KDZ1-specific probes on all ten isolates that had undergone two rounds of murine passage, as well as the cockatoo guano and PU isolates. Restriction enzymes that cut once within the TE sequence, outside of the probe site, were used to produce a range of fragment lengths (Fig S9A). The same hybridization band patterns were observed in the original cockatoo guano isolate and in all isolates recovered following infection (Fig S9B, C), indicating that no CNEST or KDZ1 mobilization occurred during infection, despite detection of CNEST transposition in vitro at 37°C (Fig 5B). Thus, while heat-stress activation of CNEST appears conserved between environmental and clinical isolates, no transposition events were detected in isolates recovered from human or murine infections.

### Full-length copies of TEs are present across multiple isolates and lineages

Because TE mobility was highly isolate- and strain-dependent in the case 15, 77, and 7 isolates, where the TEs captured mobilizing were unique to each case, we hypothesized that this may be due to the absence of full-length copies in certain isolate backgrounds. To evaluate the prevalence of full-length TE copies across *C. neoformans* isolates, we conducted homology searches in assembled genomes using the active TE sequences that were identified inserting into the *FRR1* gene as queries. Full-length copies were defined as hits with ≥ 90% nucleotide identity and ≥ 95% coverage of the query sequence. We identified full-length copies of CNIRT6, CNEST, TCN3, KDZ1, and KDZ3 in several genomes of isolates where transposition was not detected using the transposon trap assay (Fig S10A). For example, at least one full-length copy of CNEST was found in all genomes assessed, including strains from the VNI, VNII, and VNBII lineages (Fig S10A). Additionally, while Gypsy-1_CaGr was not observed mobilizing in vitro, at least one full-length copy was present in all genomes, with up to 12 copies detected in the case 7 isolates (Fig S10A). While it appears that there may be a link between increased genomic copy number and TE mobility, we do not have sufficient data to formally assess this.

To assess whether full-length copies may be present in the 15 *C. neoformans* cases initially screened without available whole-genome assemblies, we performed TE copy number estimation by aligning short reads to representative full-length TE sequences obtained from an isolate in which the TE family was identified as active. Copy numbers were estimated based on normalized read depth, defined as the depth at which ≥ 95% of the TE sequence is covered [37]. This analysis revealed that full-length copies of potentially active elements may be present even in isolates in which no TE mobilization was detected in the preliminary screen, including cases where estimated copy numbers exceeded those in isolates with observed mobilization (Fig S10B).

### DNA methylation status may explain differential mobility of TCN3

Although most TEs identified in this study were detected mobilizing exclusively in a single isolate background, additional full-length copies were found in other assemblies or inferred through copy number estimation (Fig S10A, B). This suggests that TE mobility may be differentially regulated across isolate backgrounds. For example, elevated TE mobility in some isolates may be explained by a loss of RNAi functionality or the absence of DNA methylation on TE sequences.

To explore potential regulatory mechanisms, we first assessed the levels of DNA methylation in full-length TE copies in which mobilization was either observed or not observed using the transposon trap assay. Full-length TCN3 copies were highly methylated in cases 15 and 1, whereas hypomethylated copies were observed in case 7 isolates (Table S4). In contrast, DNA methylation was generally absent from full-length DNA transposons in strains where no mobilization was observed. For example, in whole-genome assemblies, full-length KDZ1, KDZ3, and CNEST elements were unmethylated in isolates where transposition was not observed in *FRR1*, with the exception of CNEST in case 1 (Table S4). These findings suggest that while DNA methylation may play a role in regulating TCN3 mobility at the centromeres, the methylation status of full-length DNA transposons (which are primarily unmethylated) does not explain isolate- or strain-specific differences in mobility.

Next, we examined whether genes required for RNAi function were intact in cases where TE mobility was observed. In cases 7, 15, 77, and the cockatoo guano isolate, we first screened for predicted loss-of-function mutations defined as predicted protein sequences < 85% of the expected length using a previously published list of genes essential for RNAi [37]. No genes required for RNAi function exhibited truncated protein sequences. Second, we performed variant analysis and identified zero nonsense or frameshift mutations across the entire coding sequence for RNAi genes in all isolates, providing no strong evidence for loss-of-function mutations.

### DNA transposons are predominantly unmethylated and commonly found in gene-rich regions

We observed that all full-length, active DNA transposons mapped in the case 15, 77, and 7 genomes were located in gene-rich regions along the chromosome arms (Fig 2D, 3D, 4C). This pattern contrasts with the distribution of retrotransposons in *Cryptococcus*, as previous studies have shown that LTR elements predominantly localized in centromeres [42], and non-LTR elements are enriched at chromosome ends [38]. To further investigate the genomic distribution of DNA transposons in *C. neoformans*, which has not previously been described, we took a comprehensive, unbiased approach to assess genome-wide TE content and distribution. De novo TE annotation was performed using the Earl Grey package [79] for cases 1, 7, 15, and 77, as well as the genome of the reference strain H99, in which TE mobilization has not been observed unless RNAi is disrupted [38,49]. Results from the first assemblies of serially collected isolates are shown, as assemblies of later relapse isolates from the same patients yielded similar results. TEs occupied approximately 3 – 5% of the genomes analyzed (Fig 6A). While the proportion of LTR retrotransposons remained relatively consistent (2.2 – 2.6%), greater (more than tenfold) variation was observed in the proportion of DNA transposons (0.08 – 0.85%), and non-LTR retrotransposons (0.06 – 2.06%) (Fig 6A). Notably, the *C. neoformans* reference strain H99 had the lowest proportion of DNA transposons among all isolates analyzed.

**Fig 6.**
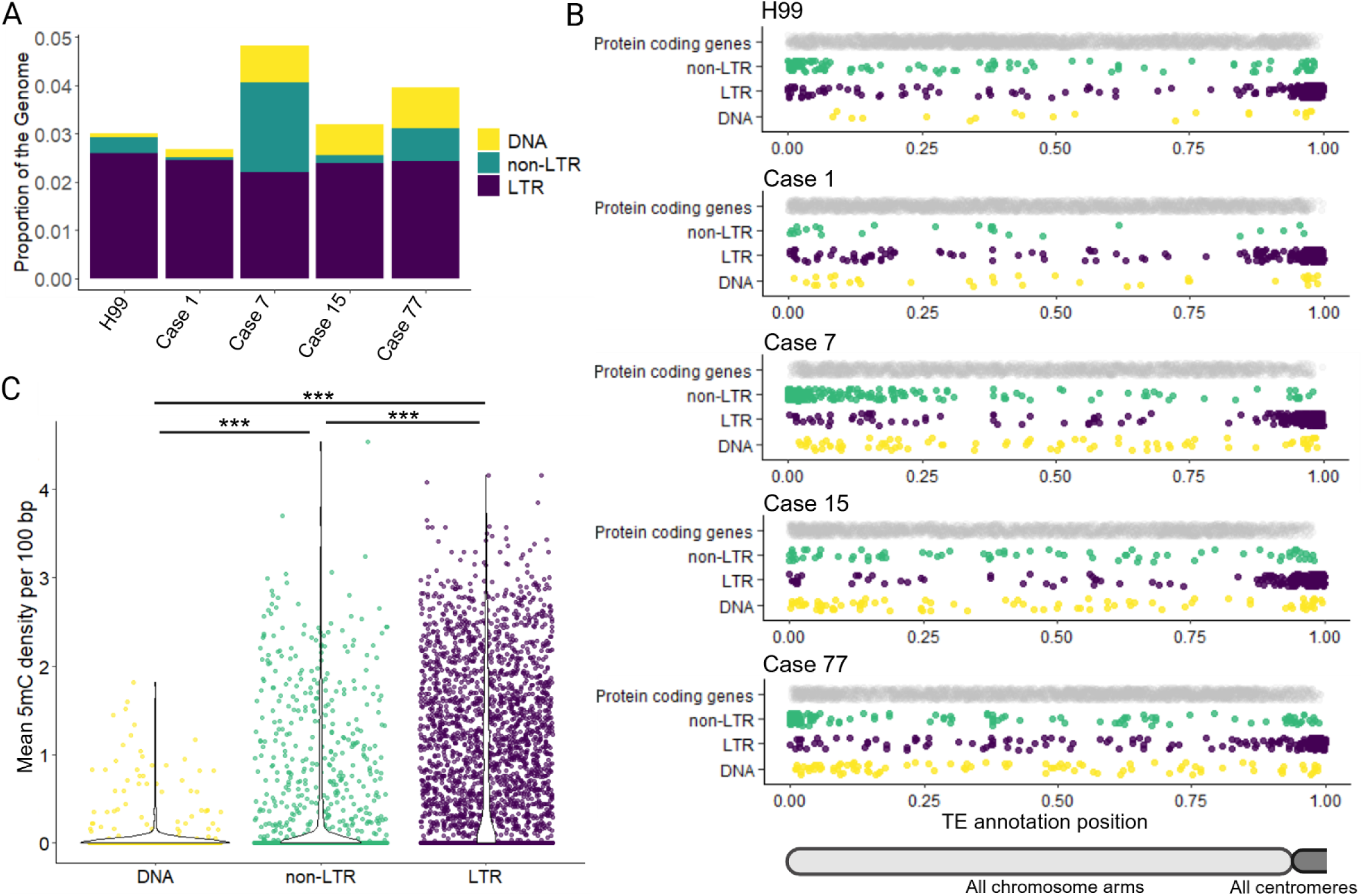
Genomic distribution and DNA methylation differ among non-LTR, LTR, and DNA transposons. (**A**) The proportion of the genome composed of non-LTR, LTR, and DNA transposons, based on Earl Grey TE annotations. (**B**) Scaled positions of protein coding genes and non-LTR, LTR, and DNA transposons across chromosome arms. Chromosomes were split at the midpoint of the centromere region, and protein-coding gene and TE positions were scaled from 0 to 1 based on chromosome arm length. Data for all chromosome arms is plotted. (**C**) Mean 5mC DNA methylation density per 100 bp for DNA, non-LTR, and LTR TEs in isolates from cases 1, 7, 15, and 77. Violin plots are overlaid to illustrate the distribution. Significance was assessed using the pairwise Wilcoxon rank-sum test with continuity correction. *** denotes Bonferroni-adjusted *p* < 2e-16. The first genome assembly from serially collected clinical isolates was chosen as a representative for all cases in (**A**), (**B**), and (**C**).

To visualize TE distributions, chromosomes were split at the midpoint of the predicted centromere. For each chromosome arm, the length, TE positions, and protein-coding gene annotation positions were scaled from 0 to 1, allowing data from all chromosome arms to be plotted on a single track. In agreement with previously published data, LTR and non-LTR elements cluster in the predicted centromeres and subtelomeres, respectively; these are regions with a lower density of protein-coding genes (Fig 6B). By contrast, DNA transposons appear to be more evenly distributed across chromosome arms (Fig 6B). We used quantile-quantile (QQ) plots to assess whether the positions of DNA transposons along chromosome arms follow a uniform distribution. For each genome, we generated theoretical quantiles as sorted random numbers between 0 and 1; the total number of theoretical quantiles matched the total number of DNA transposon annotations. The plotted points fell approximately along the diagonal of the QQ plots, indicating that DNA transposons were distributed roughly uniformly across the genome (Fig S11).

To assess how frequently DNA transposons were methylated, given their presence in gene-rich regions that are not typically methylated in *C. neoformans* [52,53], we compared their 5mC DNA methylation levels to those of LTR and non-LTR elements. First, we assessed average methylation density per 100 bp. DNA transposons exhibited significantly lower average 5mC densities compared to both non-LTR and LTR retrotransposons (pairwise Wilcoxon rank-sum test with continuity correction; adjusted *p* values < 2e-16 for both comparisons) (Fig 6C).

Additionally, LTR elements exhibited significantly higher DNA methylation densities compared to non-LTR elements (pairwise Wilcoxon rank-sum test with continuity correction; adjusted *p* value < 2e-16). Second, we assessed the proportion of TE annotations per genome containing at least one high-confidence 5mC call (methylated frequency from nanopore long-read signal data > 0.75). According to this analysis, we found that LTR elements are by far the most consistently methylated (55.6 – 71.2% of LTR retrotransposons contained at least one predicted 5mC), followed by non-LTR elements (7.7 – 53.4%), and DNA transposons (5.4 – 15.5%) (Fig S12A, B). The number of methylated LTR and non-LTR retrotransposons was significantly enriched compared to DNA transposons (Fisher’s method combined *p* values: 3.2e-64 for LTR elements and 9.2e-19 for non-LTR elements).

To estimate background levels of DNA methylation in protein-coding genes, we lifted over gene annotations from the H99 reference genome. The percentage of protein-coding genes with at least one high-confidence 5mC call ranged from 0.5 – 4.5%, indicating that DNA TEs remain significantly enriched for DNA methylation relative to endogenous protein-coding genes (Fisher’s method combined *p* value 3.9e-10), albeit at a lower frequency than LTR and non-LTR elements. Thus, the majority of DNA transposons across all genomes analyzed are unmethylated, and those that are methylated exhibit a lower methylation density compared to non-LTR and LTR retrotransposons (Fig 6C, Fig S12A, B). This suggests that DNA methylation is unlikely to play a major role in modulating the activity of DNA transposons in *C. neoformans*.

## DISCUSSION

Understanding the interplay between mobile genetic elements, environmental stressors, epigenetics, and genetic background is critical for uncovering factors that influence evolutionary rates and the capacity for adaptive change, particularly in the context of fungal pathogens. In this study, we screened a collection of clinical *Cryptococcus* isolates for transposable element (TE) insertions in the *FRR1* reporter gene and profiled rates of rapamycin and FK506 (rap+FK506) resistance, as well as mutation spectra, at both 30°C and the heat-stress condition of 37°C. We also generated telomere-to-telomere genome assemblies to assess TE copy number, genomic position, and DNA methylation status in isolates from patients with recurrent cryptococcal infections.

### *C. neoformans* hosts diverse active transposable elements

We identified seven TE families mobilizing to confer rap+FK506 resistance, including both retrotransposons (LTR and non-LTR) and DNA transposons (Fig 1). Several of these elements (TCN3, CNIRT6, KDZ3) were not previously recognized as active. In addition, we discovered a novel, active element, which we named CNEST. The diversity of active TEs in *C. neoformans* stands in striking contrast to other yeasts commonly used to study TEs, such as *Saccharomyces cerevisiae* and *Schizosaccharomyces pombe*, which harbor only LTR retrotransposons [80,81]. In these species, TE mobilization studies typically rely on TE constructs driven by inducible heterologous promoters, due in part to the low de novo transposition rates of endogenous copies [82–85]. By contrast, in *C. neoformans*, selecting for mutations at a single genomic locus frequently revealed insertions of endogenous TE copies with diverse transposition mechanisms (Figs 2 – 5). The ability to readily capture multiple active TE families mobilizing within a single *C. neoformans* strain underscores the value of this genetically tractable model for studying TE mobilization strategies and regulation in a basidiomycete.

Phylogenetic analysis of transposases from CNIRT6 and CNEST revealed that they belong to the CMC supergroup first described by Yuan and Wessler (2011) (Fig S3). However, CNIRT6 and CNEST are only distantly related and fall within two monophyletic clades among CMC elements: one containing CNIRT6 and transposases from plants, animals, and fungi, and a second containing CNEST and transposases from animals and fungi. Thus, *C. neoformans* hosts active members of two different CMC superfamilies, a notable finding given that this group of DNA transposons remains largely understudied in fungi and animals.

The use of a single reporter gene to capture TE insertions may have limited our potential for active mobile element discovery, since some TEs have preferences for specific insertion sequences or genetic compartments [11]. However, the *FRR1* reporter system recovered insertions of TCN3 and CNL1, which appear to target centromeric and subtelomeric regions, respectively. Mapping TCN3 insertions in *FRR1* suggested weak sequence specificity (Fig S6). Alternatively, TCN3 insertions may be guided by non-sequence-based features, such as centromere-specific histone variants [86] or heterochromatin-associated proteins [87], or its insertion pattern may be shaped by selective pressures. It remains possible that TCN3 insertion into *FRR1* is relatively rare and that most de novo insertions occur within centromeric regions.

CNL1 copies are enriched in subtelomeric regions in case 7 isolates and other *C. neoformans* genomes [38]; however, we also recovered insertions of CNL1 within *FRR1* (Fig 4). Phylogenetic analysis revealed that CNL1 is most closely related to R2 elements, and specifically to MoTeR, a telomere-targeting element characterized in the fungal plant pathogen *Magnaporthe oryzae* [71]. Although prior studies have reported that CNL1 preferentially inserts into copies of itself within arrays near telomeric repeats, its precise sequence specificity remains undefined. Limited mapping of CNL1 insertions into *FRR1* reveled that TSDs contained the sequence ’CACCCCT,’ which resembles the *C. neoformans* telomere seed sequence (’AACCCCCT’ or similar variants), although interpretation was complicated by premature termination of target primed reverse transcription (TPRT) at different positions within the element (Fig S6). Further work is needed to clarify CNL1’s target site specificity, including the extent to which it targets telomeric repeats like MoTeR, inserts into its own sequence, or both. Along with TCN3, these findings highlight open questions about factors driving target site selection and TE distribution across genomic compartments. Intriguingly, stress may be one such factor as nutrient deprivation alters Ty5 phosphorylation, shifting its insertion profile from heterochromatic regions to more broadly distributed sites across the genome [87].

Lastly, most observed TE mobilization events were driven by apparently full-length TE copies containing detectable functional domains required for mobilization. Phylogenetic analysis revealed full-length, seemingly intact transposase sequences within mobilized DNA transposons, and functional domains required for mobilization were identified in mobilized TCN3 and CNL1. We found no evidence for the mobilization of clearly non-autonomous TE copies (significantly shortened or lacking apparent mobilization machinery). However, we did detect truncated CNL1 insertions, likely resulting from premature termination of TPRT, as well as instances of solo TCN3 LTRs, which likely arose from non-allelic recombination between LTR sequences following TCN3 insertion. This contrasts with other organisms, such as humans, plants, and *Drosophila*, where shortened non-autonomous TEs frequently transpose [88–90].

Collectively, we identified seven TE families in *C. neoformans*: DNA transposons from the KDZ superfamily and CMC supergroup, alongside a Ty3/Gypsy LTR element and an R2 non-LTR element actively transposing. These TEs employ distinct transposition mechanisms and show preferences for different genomic compartments, revealing a rich “ecosystem” of active TEs within *C. neoformans*.

### Transposable element mobilization in *C. neoformans* is strongly influenced by temperature

Previous work in *C. deneoformans* demonstrated that heat stress increases TE mobilization [35,36]. While some mobile elements in *C. neoformans* followed this pattern, mobilizing more frequently at elevated temperature, we also uncovered TEs that behave differently. Several elements (CNL1, KDZ1, KDZ3) were found to mobilize exclusively at 30°C in vitro. Others (CNIRT6, CNEST) mobilized at both temperatures, but with higher frequency at 37°C. Therefore, by limiting our initial screen to 37°C and examining only ten independent mutants, we may have overlooked strains harboring mobile elements that are active only at lower temperatures or mobilizing at low frequency. Interestingly, we found completely non-overlapping sets of mobilizing TE families in *C. neoformans* and *C. deneoformans* [35,36]. The discovery of TE families that mobilize more frequently at lower temperatures adds further complexity to this system, suggesting the involvement of multiple TE- or host-specific mechanisms in temperature-regulated mobility in *Cryptococcus*. One such mechanism may involve TCN3 exploiting the host’s heat shock response via heat shock elements in its LTR to facilitate mobilization (Fig 1E). While heat shock-induced TE activation via host machinery has been demonstrated in plants and animals [91–93], it remains undescribed in fungi.

Other temperature-dependent effects on TE mobility may result from (1) impacts on functionality of TE-encoded machinery, (2) altered TE suppression, or (3) changes in chromatin accessibility at *FRR1* facilitating transposition. First, Ty1 mobilization in *S. cerevisiae* is most frequent at 22°C and absent above 34°C due to inactivity of the Ty1-encoded protease [94].

Similarly, non-functional TE machinery may explain, for instance, why elements from the KDZ superfamily were only captured mobilizing at 30°C. Second, heat stress can disrupt TE suppression: in *Drosophila*, Hsp70 displaces PIWI-interacting RNA (piRNA) biogenesis factors to the lysosome for degradation, leading to transcriptional activation of TEs [95]. In *C. neoformans*, a recent study found a link between the Calcineurin signaling pathway, which is necessary for growth at 37°C, and RNAi-mediated silencing of a repetitive transgene, suggesting a potential mechanism for altered RNAi function during heat stress [96]. Third, heat stress may increase chromatin accessibility at *FRR1*, facilitating higher TE insertion rates independent of any direct effects on TE suppression or machinery [97,98]. Elucidating the mechanisms underlying differential TE mobilization under heat stress in *C. neoformans* will advance our understanding of the role of TEs in adaptive evolution during infection and in other heat-stress contexts. Additionally, it is likely that other environmental factors impact TE mobilization, given that RPMI medium altered TE dynamics across most strains, although it remains unclear whether this is due to changes in nutrient availability, pH, or other factors (Fig S7). Moreover, although CNL1 mobilization was only observed in vitro at 30°C, it was the sole TE implicated in genomic changes during recurrent cryptococcal infection, highlighting that temperature is not the sole regulator.

### DNA transposons are hypomethylated relative to retrotransposons, and are commonly found in gene-rich regions

Given that *C. neoformans* lacks a de novo cytosine DNA methyltransferase, DNA methylation as a TE suppression mechanism is relatively inflexible and unlikely to target newly inserted or unmethylated TE sequences [52]. Despite this, a substantial proportion of TEs exhibit DNA methylation: 55.6 – 71.2% of LTR elements and 7.7 – 53.4% of non-LTR elements (Fig S12). In contrast, DNA transposons are less frequently methylated than retrotransposons, with only 5.4 – 15.5% showing methylation. When methylated, DNA transposons exhibit significantly lower average 5-methylcytosine (5mC) density compared to LTR and non-LTR elements (Fig 6). Nonetheless, DNA TEs are still significantly enriched for DNA methylation compared to protein-coding genes. This reduced level of methylation may contribute to the more frequent in vitro mobilization of DNA transposons observed in this study, where five of the seven mobile elements identified were DNA TEs. Furthermore, full-length DNA transposons are more evenly distributed across the genome and are commonly found within gene-rich regions compared to LTR and non-LTR retrotransposons (Fig 6). As a result, DNA TEs may be more likely to mediate genetic changes that alter the expression or function of protein-coding genes.

### Transposable elements provide a mechanism for rapid genetic change and facilitate adaptation in *C. neoformans*

Growth at mammalian body temperature (37°C) significantly increased the rate of acquired resistance to rap+FK506 in both clinical and environmental isolates (Figs 2–5). In all cases, this effect was driven by elevated TE insertion rates, highlighting the role of TEs in generating adaptive phenotypes such as antifungal resistance. These findings suggest that stress may enhance evolvability within *C. neoformans* populations through bursts of rapid TE mobilization.

RNA interference (RNAi) is the primary known mechanism for TE suppression in *C. neoformans*, and loss of RNAi function is associated with hypermutator phenotypes driven by uncontrolled TE activity [38]. However, recent work has shown that this is not always the case, as some isolates with loss-of-function mutations in essential RNAi genes maintain low TE burdens and mutation rates [37]. In this study, none of the clinical or environmental isolates profiled harbored predicted loss-of-function mutations in RNAi pathway genes. Nevertheless, isolates from cases 15 and 77 exhibited rap+FK506 resistance rates at 37°C comparable to those observed in hypermutator strains, including the mismatch repair-deficient *msh2Δ* mutant [37,38]. This suggests that, in certain strain backgrounds, heat stress alone can be sufficient to induce a hypermutator-like phenotype through increased TE mobility, even in strains with a relatively low TE burden (e.g., two full-length copies of CNIRT6 in case 15 isolates).

When assessing the impact of TEs on in-host microevolution, we observed an expansion of CNL1 copy number in serial isolates collected from a patient with recurrent cryptococcal meningitis, suggesting a role for TE activity in within-host evolution (Fig 4). However, no transposition or TE-mediated genomic changes were detected in other longitudinally sampled clinical isolates or in isolates collected following murine passaging. This may suggest that TE mobilization events are rare in vivo, that TE-mediated changes did not confer a selective in-host growth advantage during infection, or that TE-mediated changes occurred in subpopulations not captured by sampling a single isolate. Previous studies in *C. deneoformans* found evidence of transposition occurring during murine infection [35,36]. In these cases, however, only drug-resistant isolates were selected, requiring that loss-of-function mutations at specific reporter loci had occurred. In contrast, there was no selective pressure for evolved drug resistance or other traits during the recovery of patient and mice isolates examined in this study.

### Genetic background influences TE mobility

Telomere-to-telomere genome assemblies revealed that the presence of full-length TEs does not reliably predict their mobilization in vitro. For instance, in clonally related case 7 isolates, we observed striking differences in KDZ2 and KDZ3 mobilization at 30°C, despite these elements being present at identical copy number and genomic positions, with identical sequences, and lacking DNA methylation in both isolates. Interestingly, of the two single nucleotide polymorphisms (SNPs) previously identified between case 7 isolates, one was an amino acid substitution in *SET5* (CNAG_03890), a predicted protein lysine methyltransferase [29]. This suggests that alternative epigenetic regulation may be driving differential mobility.

Similarly, unmethylated full-length CNEST copies were identified in all assembled genomes except for case 1; however, mobilization was observed only in the case 77 and cockatoo guano isolates. Notably, these isolates had the highest CNEST copy numbers, suggesting that copy number may be positively correlated with increased mobility in vitro. These observations suggest that isolate- and strain-specific TE mobility may be regulated by TE copy number polymorphisms, sequence variation within TEs, differential targeting by RNAi, or genetic variation in host factors that control transposition.

Altogether, our findings highlight the complex interplay between genetic background, epigenetics, and environmental conditions in governing TE mobilization, with temperature emerging as a major determinant. Further studies should investigate the extent to which stress-induced transposition generates genetic diversity and whether this diversification drives adaptation to infection-relevant stressors and the repeated emergence of antifungal drug resistance in *C. neoformans*.

## METHODS

### Strains and growth media

The strains used in this study are listed in Table S1. Strains were grown in YPAD medium (1% yeast extract, 2% peptone, 2% dextrose, 0.025% adenine) with 2% agar added for solid media. For host-mimicking conditions, strains were cultured on RPMI plates containing 3.45% MOPS, 0.2% glucose, 2% agar, and 1.04% RPMI 1640 powder (Sigma #R1383). Mutants resistant to rapamycin and FK506 (rap+FK506) were selected for on YPAD plates supplemented with 1 μg/mL rapamycin (LC Laboratories) and 1 μg/mL FK506 (Astellas).

### Fluctuation assay and mutation spectra

Cultures were inoculated with independent colonies. Each colony was split into 1 mL cultures of YPAD and grown rotating at 30°C or 37°C for 3 days, allowing all cultures to reach saturation. Cells were pelleted, washed, and resuspended in water. Cells were appropriately diluted, plated on solid YPAD medium, and incubated at 30°C for 3 days. Colony counts were used to calculate colony forming units (CFUs) per culture. The remainder of the culture was plated on two rap+FK506 plates and incubated at 37°C for four days before counting colonies. Total CFUs and rap+FK506-resistant colony counts were used to independently calculate the drug resistance rate for each isolate and temperature using the Fluctuation AnaLysis CalculatOR (FALCOR), applying the Ma-Sandri-Sarkar (MSS) maximum likelihood method [99]. For all fluctuation assays a minimum of 23 cultures were used to calculate the drug resistance rate. A single rap+FK506-resistant colony from each culture was selected as an independent mutant for genomic DNA isolation, performed using either in-house reagents or a MasterPure Yeast DNA Purification Kit (VWR #76081-694). A minimum of 24 independent drug-resistant mutants were collected per isolate and temperature for the YPAD mutation spectra analysis.

The *FRR1* gene was amplified for Sanger (Azenta) or linear amplicon long-read sequencing (Plasmidsaurus). Amplification of *FRR1* was carried out using a variety of polymerases: MyTaq Mix (Thomas Scientific #C755G91), Apex Taq RED Master Mix (Genesee Scientific #42-138), Platinum SuperFi II PCR Master Mix (ThermoFisher #12368010), Q5 High-Fidelity DNA Polymerase (NEB #M0491S), and LongAmp Taq DNA Polymerase (NEB #M0323S). We often observed “phantom” wild-type–sized secondary bands when amplifying across TEs, which can occur as a PCR artifact when amplifying across long repetitive DNA sequences [100]. In these instances, gel extraction using the Zymoclean Gel DNA Recovery Kit (Zymo Research #D4007) was performed on the higher molecular weight bands, which were then sent for sequencing. For TEs identified as inserting into *FRR1*, junction PCR with primers targeting the internal TE sequence and primers flanking *FRR1* were used to confirm a subset of insertion events, as PCR amplification across long repetitive TE sequences can be difficult [100]. Primers used to amplify *FRR1* and for TE-specific junction PCR are listed in Table S5.

Sanger sequencing data were analyzed in SeqMan Ultra 17 or DNASTAR Navigator 17 by aligning reads to the wild-type *FRR1* sequence for each isolate. TEs were assigned to families based on the 80-80-80 rule: nucleotide similarity greater than 80%, over more than 80 base pairs, covering at least 80% of the recovered sequence length [40]. The TE consensus sequence for TCN3 was obtained from Repbase. For TEs lacking family-level consensus sequences, GenBank submissions were used to assign family membership (GenBank accessions: PQ181658 [KDZ1], PQ181660 [KDZ2], PQ181661 [KDZ3], and JQ277267.1 [CNIRT6]). CNL1 family membership was called based on sequence similarity to CNL1 mapped by Priest et al. (2022) in Bt65 and Bt81 genome assemblies (BioProject accession PRJNA749953). Domains were identified within full-length TE sequences using InterPro [101].

### RPMI mutation spectra

To generate independent mutants during growth on RPMI medium, a prong-plating procedure was carried out. Single colonies were inoculated in liquid YPAD and grown overnight rotating at 30°C, then cells were washed and diluted in water. A custom tool containing 121 flat-tipped pins was used to transfer approximately ∼10^4^ cells/pin onto non-selective RPMI plates [102]. To confirm there were no pre-existing mutations, cells were also spotted on rap+FK506 plates. Confluent spots of cells on these plates would indicate mutations occurred during the overnight growth in YPAD and prong plating would not be possible to generate independent mutants. If less than 5 colonies appeared on these control plates, experiments were continued. RPMI plates were incubated for 3 days at either 30°C or 37°C. RPMI plates were then replica plated to rap+FK506 plates and incubated at 37°C for 3 days. A single colony was picked from each spot for genomic DNA isolation, followed by *FRR1* amplification and either Sanger sequencing (Azenta) or junction PCR. A minimum of 18 independent drug-resistant mutants were collected per isolate and temperature for the RPMI mutation spectra analysis. *FRR1* amplification, junction PCR, Sanger sequencing data analysis, and TE classification were carried out as described above.

### TE insertion rate

TE insertion rate was calculated using the proportion of independent mutants with a given TE insertion multiplied by the drug resistance rate. The 95% confidence intervals (CIs) for proportions were determined using the vassarstats.net proportion CI calculator including continuity correction. 95% CI for TE insertion rates were determined by a method employing the square root of the sum of the squares (using the component 95% CI for rates and proportions) [103].

### Transposase and reverse transcriptase protein sequence alignments and phylogenies

Transposase and reverse transcriptase protein sequences were obtained from Yuan and Wessler (2011) for CMC (CACTA, Mirage, Chapaev) related sequences, Repbase consensus sequences (https://www.girinst.org/), and from BLASTp (for translated mRNA sequences) or tBLASTn (for translated genomic sequences) via NCBI (https://blast.ncbi.nlm.nih.gov/Blast.cgi). Protein sequences were aligned using MAFFT (https://www.ebi.ac.uk/jdispatcher/msa/mafft) with the following parameters, protein matrix = BLOSUM62, gap opening penalty = 1.53, gap extension penalty = 0.123, tree rebulding number = 2, max iterations = 2, and FFTS = none. Aligned protein sequences were then used to construct maximum likelihood phylogenies with 100 bootstrap replicates using MEGA version 11 [104].

### Nanopore sequencing

To extract high molecular weight DNA for library preparation, overnight cultures were grown in 50 mL liquid YPAD shaking at 30°C. Cell were pelleted and lyophilized overnight. Genomic DNA was extracted using a cetyltrimethylammonium bromide (CTAB) extraction method with some modifications [105]. Briefly, cells were lysed by bead beating, and DNA extraction buffer (100 mM Tris-HCl, pH 7.5; 1% w/v CTAB; 0.7 M NaCl; 10 mM EDTA; 1% v/v 2-mercaptoethanol) was added and incubated at 65°C for 30 minutes. Chloroform purification was then performed, and layers were separated by centrifugation. The aqueous layer was isolated, and DNA was subsequently precipitated, washed, and treated with RNase A. Sodium acetate (NaOAc) was then added to a final concentration of 0.6 M, and chloroform purification was repeated. DNA was precipitated again, washed, and spooled to collect high molecular weight DNA.

DNA quality was assessed using NanoDrop One A260/A280 and A260/A230 ratios (Thermo Fisher Scientific) and quantified using Qubit dsDNA Broad Range Assay Kit (Invitrogen #Q32850) and the Qubit 4 Fluorometer (Invitrogen). To assess DNA fragment length, clamped homogeneous electric field (CHEF) gel electrophoresis was performed using the CHEF Mapper (Bio-Rad). Libraries were prepared using the SQK-LSK110 or SQK-NBD114.24 and EXP-NBD104 kits and sequenced using the MinION Mk1C following manufacturer’s instructions (Oxford Nanopore). Sequencing was performed using the MinKNOW software at the default voltage for at least 48 hours. Basecalling was performed using the latest Guppy software (v7.1.4, v6.3.8, or v6.4.8), or with Dorado (v0.8.1). The specific sequencing kit, flow cell, and basecalling methodology used for each isolate is listed in Table S3.

### Genome assembly

Demultiplexing was performed using Guppy (v6.5.7). The Canu (v2.2) genome assembler was used to assemble long reads into contigs, small contigs likely representing mtDNA or contaminants (< 200 kb) were discarded [106]. For the case 15 incident isolate only, assembly was performed by combining outputs from Canu and Flye (v2.9.5) [107]. Genome polishing was performed using long reads generated in this study and short reads available under the SRA umbrella project PRJNA37160 [29]. Long reads were mapped to contigs using Minimap2 (v2.24) followed by Medaka (v1.5.0 or v2.0.1) polishing using the appropriate model [108]. Short reads were mapped to the draft assembly using Bowtie2 (v2.4.4) followed by Pilon (v1.24) polishing, which was performed in five iterations, re-mapping the short reads to each new draft assembly before the next iteration of polishing [109,110]. Canu-corrected Nanopore reads were mapped to the assembly using Minimap2 (v2.24) and coverage was visually inspected in the Integrative Genome Viewer (IGV) [111]. To confirm generation of telomere-to-telomere assemblies, contig ends were inspected for the presence of telomere seed sequences [112]. Assembly breaks at the rDNA locus were artificially merged and separated by 50 Ns. Contigs were oriented to match the first serially collected isolate assembly to produce final nuclear genome assemblies. For each isolate, Canu parameters, polishing specifications, and assembly statistics are provided in Table S3.

### DNA methylation detection

5-methylcytosine (5mC) DNA methylation at CpG sites was called using Nanopolish (v0.14.0) or f5c (v1.5) (see Table S3 for isolate-specific details). Minimap2 (v2.24) was used to map long reads to the final nuclear assembly for each isolate. For DNA methylation detection using f5c, POD5 files were converted to BLOW5 using blue-crab (v0.2.0). The Nanopolish helper script methylation_frequency.py was used to calculate the methylation frequency for CpG sites.

### Centromere mapping

Predicted centromere regions were determined using a synteny based approach using the H99 reference genome (GCA_000149245.3) where centromere regions have been experimentally identified [38,42,112]. The 5 kb regions flanking H99 centromeres were used as BLASTn (v2.14.0+) queries against assembled genomes for centromere mapping [113,114].

### Full-length TE mapping in assembled genomes

To estimate full-length TE copy numbers from assembled genomes BLAT (v35) and BLASTn (v2.14.0+) were used, full-length TE sequences obtained from linear amplicon long-read sequencing of TE insertions into *FRR1* were provided as queries [59]. Full-length copies were defined as hits with identity ≥ 90% and alignment length ≥ 95% of the query length. For BLAT the -minIdentity=90 specification was used, and pslReps (v469) filtered hits to find the best alignment with a minimum coverage of 95% (-minCover=0.95). BLASTn hits meeting full-length criteria were filtered in R (v4.3.3). In instances were BLAT and BLASTn reported different full-length copy number estimates, the higher estimate was used. Only BLASTn was used to detect copies of CNL1 because BLAT reported overlapping CNL1 copies leading to overestimation, this was not observed for other TEs.

### Identification of structural rearrangements and synteny analysis

Whole-genome alignment was performed using Minimap2 (v2.24). Dotplots to visualize primary alignments between assemblies were generated in R (v4.3.3) using pafr (v0.0.2). Synteny and Rearrangement Identifier (SyRI) (v1.6.3) was used to define syntenic regions and structural rearrangements between serial isolates using default parameters [61]. Plotsr was used to visualize syntenic regions [115].

### Southern analysis

Genomic DNA was prepared using a CTAB DNA extraction method (described above) and digested with restriction enzymes BspEI, PvuII, or PspXI for CNIRT6, MLDT, and KDZ1, respectively (New England Biolabs) [105]. Digests were run on 0.7% agarose gels at 4°C for approximately 16 h. Fragments from the gel were transferred to a charged nylon membrane (Roche). Target probes were prepared using chemiluminescent DIG-labeling (Roche), see Table S5 for primers used to amplify probes. Hybridization of the DIG probes (DIG Easy Hyb Granules, Roche) and detection by chemiluminescence using an anti-DIG antibody (Anti-Digoxigenin-AP Fab Fragments, Roche) were performed according to the recommended manufacturer’s protocols and imaged on film (Amersham Hyperfilm ECL).

### TE annotation and repeat masking

Transposon annotation was performed using Earl Grey (v4.1.0) with default parameters [79]. Insertions, deletions, and duplications identified by SyRI were repeat masked using Genetic Information Research Institute (GIRI) Repbase CENSOR [78].

### Gene annotation

Gene annotations were lifted over from the H99 reference genome (GCA_000149245.3) by Liftoff (v1.6.3) with the -polish flag [116].

### TE copy number estimation using short reads

TE copy number estimation using short reads was performed as previously described [37]. Briefly, publicly available short reads (PRJNA37160) were mapped to the H99 reference genome (GCA_000149245.3) using Minimap2 (v2.24), mosdepth (v0.3.10) was used to calculate the median read depth in 500 bp non-overlapping windows, the median depth reported was used for normalization [117]. Short reads were also mapped using Minimap2 to a multi-fasta containing representative full-length TE sequences identified inserting into *FRR1*. Mosdepth was then used to produce the cumulative read depth distribution. For each TE, read depth at 0.95 coverage was normalized by the genome-wide median read depth to estimate full-length copy number.

### RNAi gene loss-of-function prediction

Genome feature annotation was “lifted over” from the H99 reference genome (FungiDB R68) to each isolate-specific genome assembly using the software tool Liftoff (v1.6.3) [116]. The ‘-polish’ option of Liftoff was employed to re-align exons in cases where the lift-over procedure resulted in start/stop codon loss or introduced an in-frame stop codon. Based on the polished lift-over annotation, the AGAT GTF/GFF Toolkit software (https://github.com/NBISweden/AGAT) was used to predict protein sequences for all annotated genes in each strain-specific assembly using the ‘agat_sp_extract_sequences.pl’ script. Where multiple protein isoforms are annotated in the reference genome, predictions were generated for each isoform. For each RNAi-related gene of interest, the predicted amino acid sequences for each isolate were compared to the H99 reference genome. Candidate loss-of-function alleles were classified as those encoding proteins whose predicted length is < 85% of the length in H99.

Variant analysis of RNAi genes was performed by aligning the coding sequence of each RNAi gene from each isolate’s polished genome assembly using the corresponding Liftoff gene annotation to the H99 reference genome coding sequence using MAFFT (v7.526) [118]. Snp-sites (v2.5.1) was then used to generate a variant call file, and the impact of variants was assessed using SnpEff (v5.3a) [119,120].

## DATA AVAILABILITY

Raw nanopore sequencing reads and whole-genome assemblies generated from isolates used in this study are available on NCBI’s Sequence Read Archive (SRA) under BioProject PRJNA1222431, accession numbers are listed in Table S3. Full-length TE sequences identified inserting into *FRR1* were uploaded to GenBank, accession numbers are listed in Table S2.

## ACKNOWLEDGEMENTS

We thank Dr. Marco Dias Coelho for training and assistance with de novo genome assembly. We also thank Dr. Arturo Casadevall for providing the cockatoo guano isolate, as well as the isolates collected after murine passage. We thank Drs. Joseph Heitman and Jun Huang for helpful discussions and for sharing unpublished results on KDZ transposons in *C. neoformans*. Finally, we thank Dr. Sue Jinks-Robertson for her advice and discussions during project conceptualization. The isolates from patients with recurrent cryptococcal meningitis in this study were collected through GERMS-SA surveillance, provided by Dr. N.P. Govender of the National Institute for Communicable Diseases, South Africa, and shared by Dr. J. Perfect at Duke University. We thank members of GERMS-SA, 2005 to 2008, including S. Vasaikar (Eastern Cape); N. Janse van Rensberg, A. Möller, P. Smith, and A. M. Pretorius (Free State); K. Ahmed, A. Hoosen, R. Lekalakala, P. P. Sein, C. Feldman, A. S. Karstaedt, O. Perovic, J. Wadula, M. Dove, K. Lindeque, L. Meyer, and G. Weldhagen (Gauteng); S. Harvey and P. Jooste (Northern Cape); D. Cilliers and A. Rampe (North West Province); W. Sturm, T. Vanmali, P. Bhola, P. Moodley, S. Sithole, and H. Dawood (KwaZulu Natal); K. Hamese (Limpopo); K. Bauer, G. Hoyland, J. Lebudi, and C. Mutanda (Mpumalanga); R. Hoffmann, S. Martin, L. Liebowitz, and E. Wasserman (Western Cape); A. Whitelaw (Western Cape); A. Brink, I. Zietsman, M. Botha, X. Poswa, M. da Silva, and S. Budavari (Ampath laboratories); C. Heney and J. Smit (Lancet laboratories); M. Senekal (Pathcare laboratories); A. Schuchat and S. Schrag (CDC); K. P. Klugman (Emory); and C. Cohen, L. Dini, L. de Gouveia, J. Frean, S. Gould, K. Keddy, K. M. McCarthy, J. Patel, S. T. Meiring, E. G. Prentice, V. C. Quan, J. Ramalivhana, A. Sooka, A. von Gottberg, and N. P. Govender (National Institute for Communicable Diseases).

## COMPETING INTERESTS

The authors have declared that no competing interests exist.

## ABBREVIATIONS

5mC: 5-methylcytosine
CMC: CACTA, Mirage
Chapaev Indel: Insertion-deletion
KDZ: Kyakuja, Dileera, Zisupton
LTR: Long terminal repeat
Rap+FK506: Rapamycin and FK506
RNAi: RNA interference
SNP: Single nucleotide polymorphism
TSD: Terminal site duplication
TE: Transposable element
TIR: Terminal inverted repeat

---

**Fig S1.**
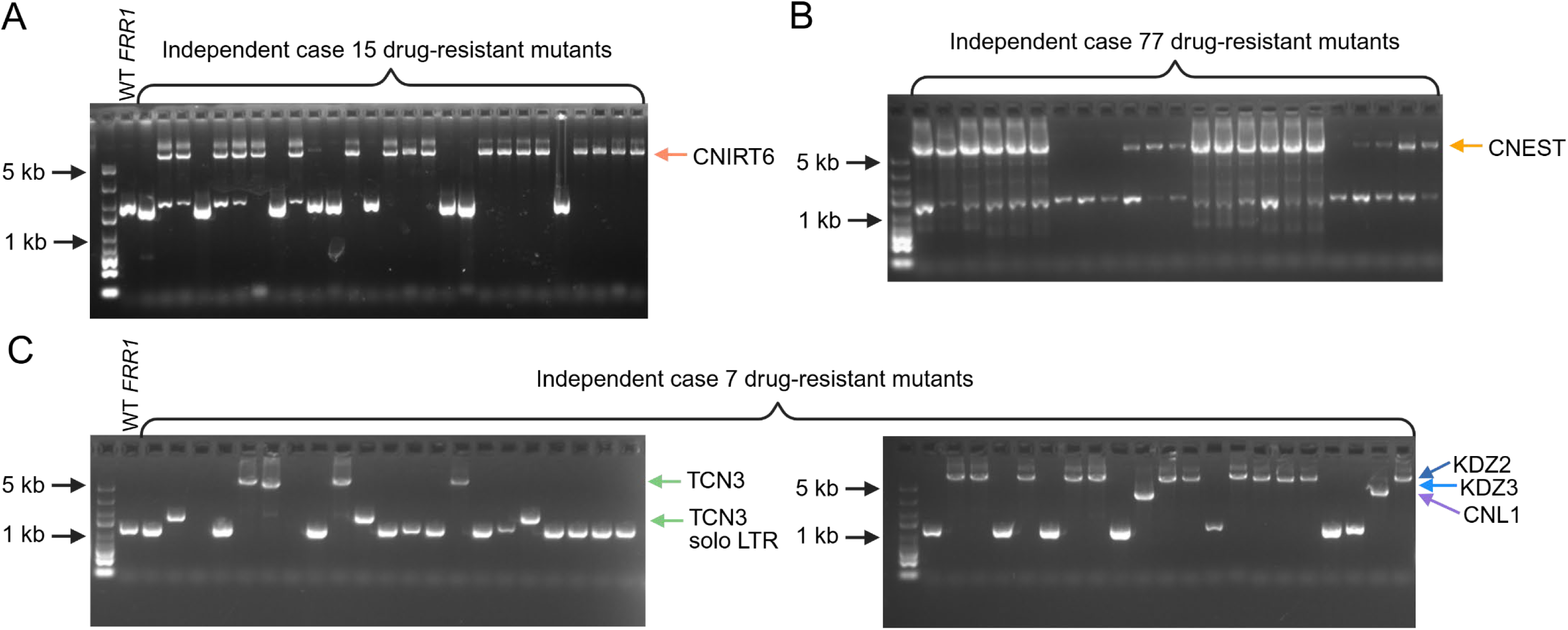
PCR amplification of *FRR1* in wild-type and independent rap+FK506-resistant mutants. *FRR1* was amplified from independent drug-resistant mutants from case 15 (**A**), case 77 (**B**), and case 7 (**C**); a subset of representative mutants are shown. Higher molecular weight bands indicative of TE insertion in *FRR1* were verified by Sanger or linear amplicon sequencing, or junction PCR. Colored arrows indicate the molecular weights of bands identified as TEs.

**Fig S2.**
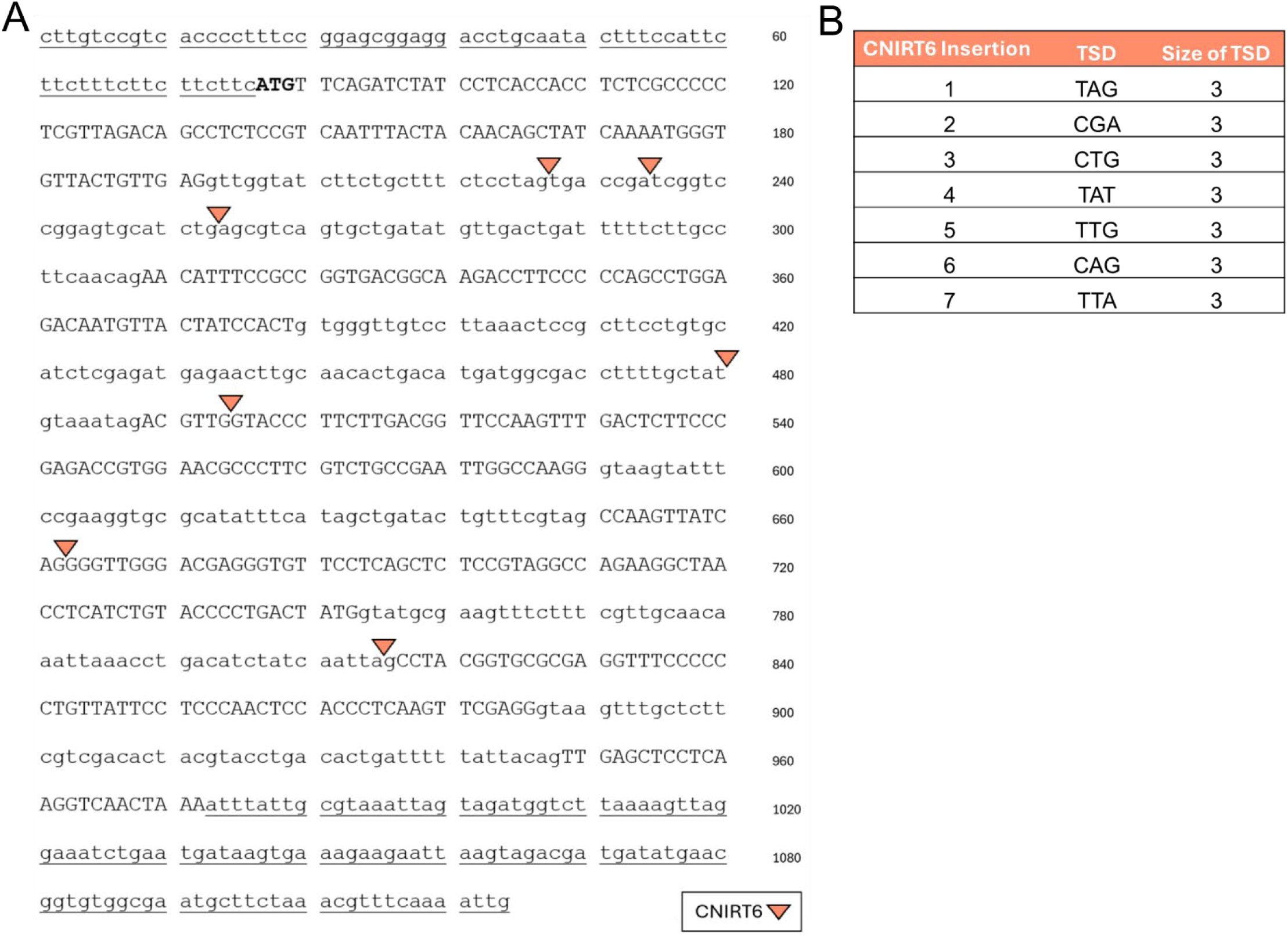
CNIRT6 insertion into *FRR1*. (**A**) A subset of CNIRT6 insertion sites (orange arrows) mapped in *FRR1*. The 5′ and 3′ untranslated regions (UTRs) of *FRR1* are underlined; introns are shown in lowercase, exons in capital letters, and the start codon is bolded. (**B**) TSDs detected at CNIRT6 insertion sites. Insertions are ordered based on their appearance in the *FRR1* sequence from beginning to end.

**Fig S3.**
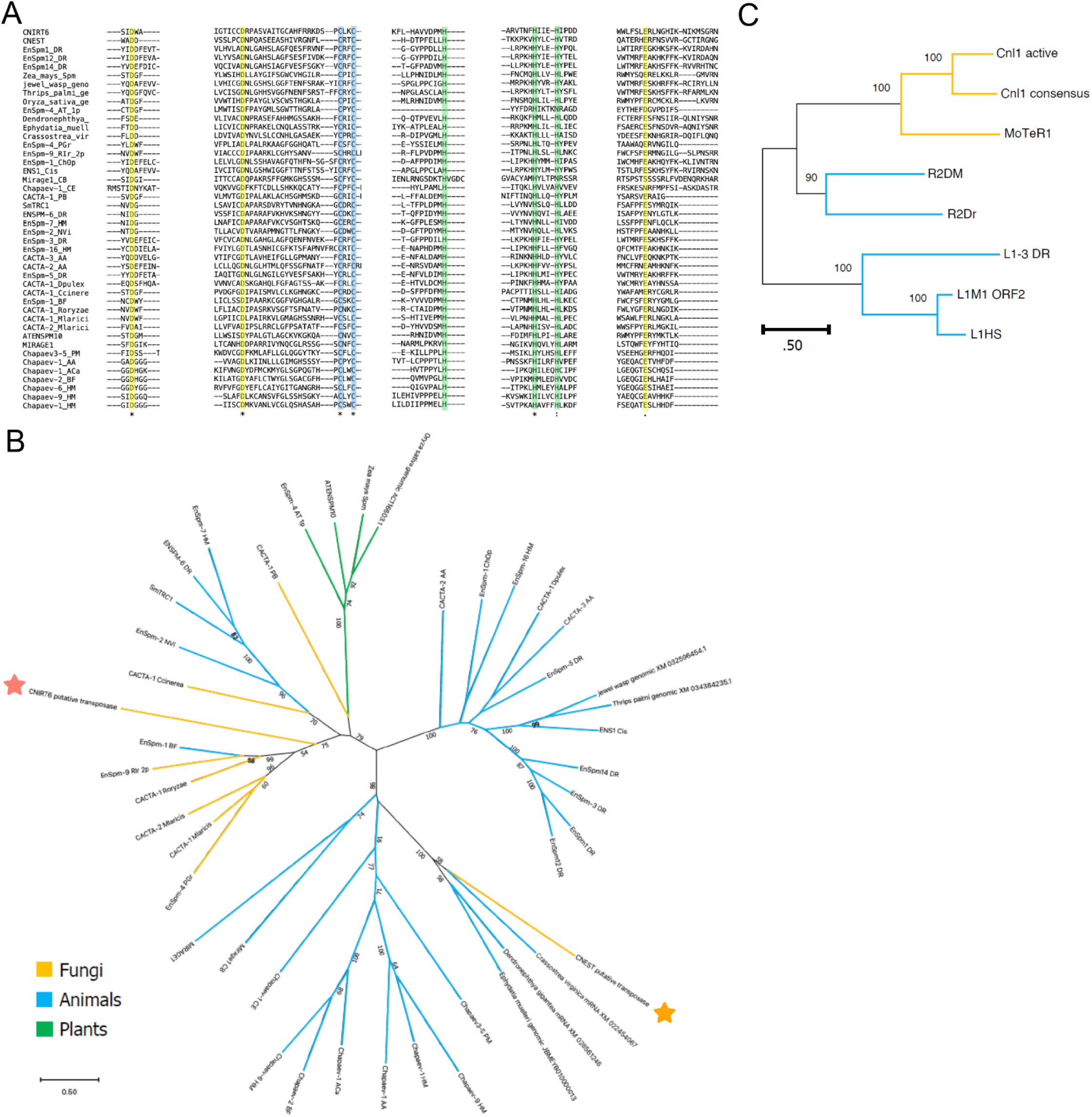
Classification of CNIRT6, CNEST, and CNL1. (**A**) Regions of interest in a multiple sequence alignment highlighting conserved residues in CMC supergroup elements, which include members of the CACTA (also known as EnSpm), Mirage, and Chapaev superfamilies. Highlighted residues include the DDE catalytic core (yellow), conserved cysteine residues (blue) and conserved histidine residues (green) [41]. (**B**) Unrooted amino acid phylogeny of diverse CMC transposases, demonstrating the relationship of CNIRT6 and CNEST with known CMC families and other related sequences. CNIRT6 is marked with an orange star, and CNEST with a gold star. (**C**) Unrooted amino acid phylogeny of reverse transcriptase domains, demonstrating the relationship between CNL1 and MoTeR1, and more distantly related R2 elements from flies and fish, as well as mammalian L1 elements. Branches in (**B**) and (**C**) are colored by kingdom: animals (blue), plants (green), and fungi (yellow). Bootstrap values greater than 50% are shown at nodes. Scale bars represent amino acid substitutions per site. Abbreviations: HS – *Homo sapiens*, DR – *Danio rerio*, DM – *Drosophila melanogaster*, CE – *Caenorhabditis elegans,* CB – *Caenorhabditis briggsae*, PM – *Petromyzon marinus*, BF – *Branchiostoma floridae*, Dpulex – *Daphnia pulex*, Cis – *Ciona intestinalis*, Aca – *Aplysia californica*, Sm – *Schistosoma mansoni*, AA – *Aedes aegypti*, ChOp – *Chionoecetes opilio*, Jewel wasp/NVi – *Nasonia vitripennis*, HM – *Hydra magnipapillata*, Ccinerea – *Coprinopsis cinerea*, RIr – *Rhizophagus irregularis*, Roryzae – *Rhizopus oryzae*, Mlaricis – *Melampsora laricis*, PGr – *Puccinia graminis*, PB – *Phycomyces blakesleeanus*, Mo – *Magnaporthe oryzae*, AT – *Arabidopsis thaliana*.

**Fig S4.**
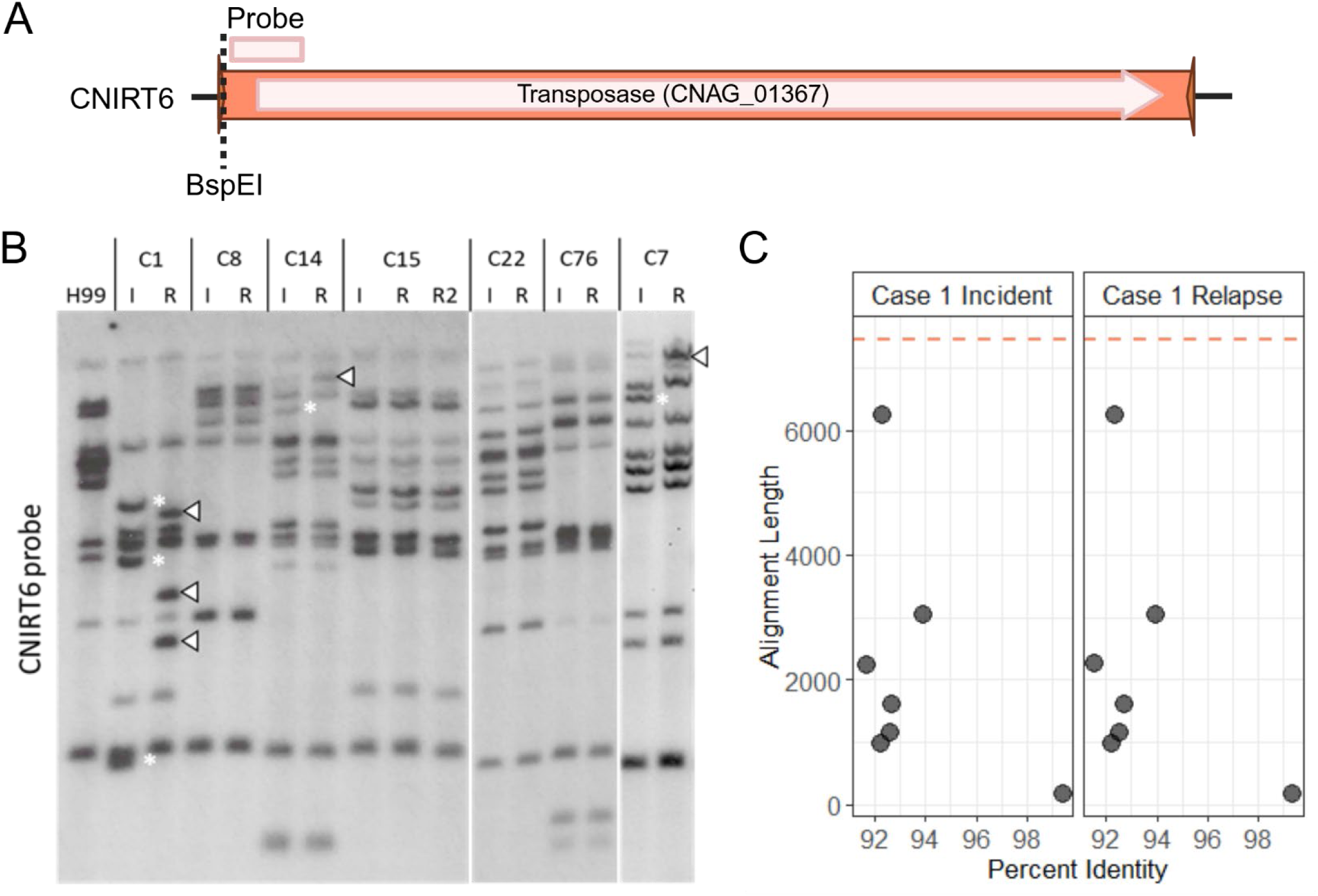
Putative CNIRT6 transposition events in serially collected clinical isolates. (**A**) Location of the restriction site (dotted line) and CNIRT6-specific probe used in Southern analysis. (**B**) Southern blot of genomic DNA digested with BspEI probed for CNIRT6 in incident (I) and relapse (R) isolates; case numbers are indicated as C#. Asterisks denote the loss of CNIRT6 fragments, while arrowheads indicate the emergence of CNIRT6 fragments at new genomic locations between incident and relapse isolates. (**C**) BLAT homology search hits identified in case 1 isolates using full-length CNIRT6 as the query. The orange dashed line indicates the length of full-length CNIRT6.

**Fig S5.**
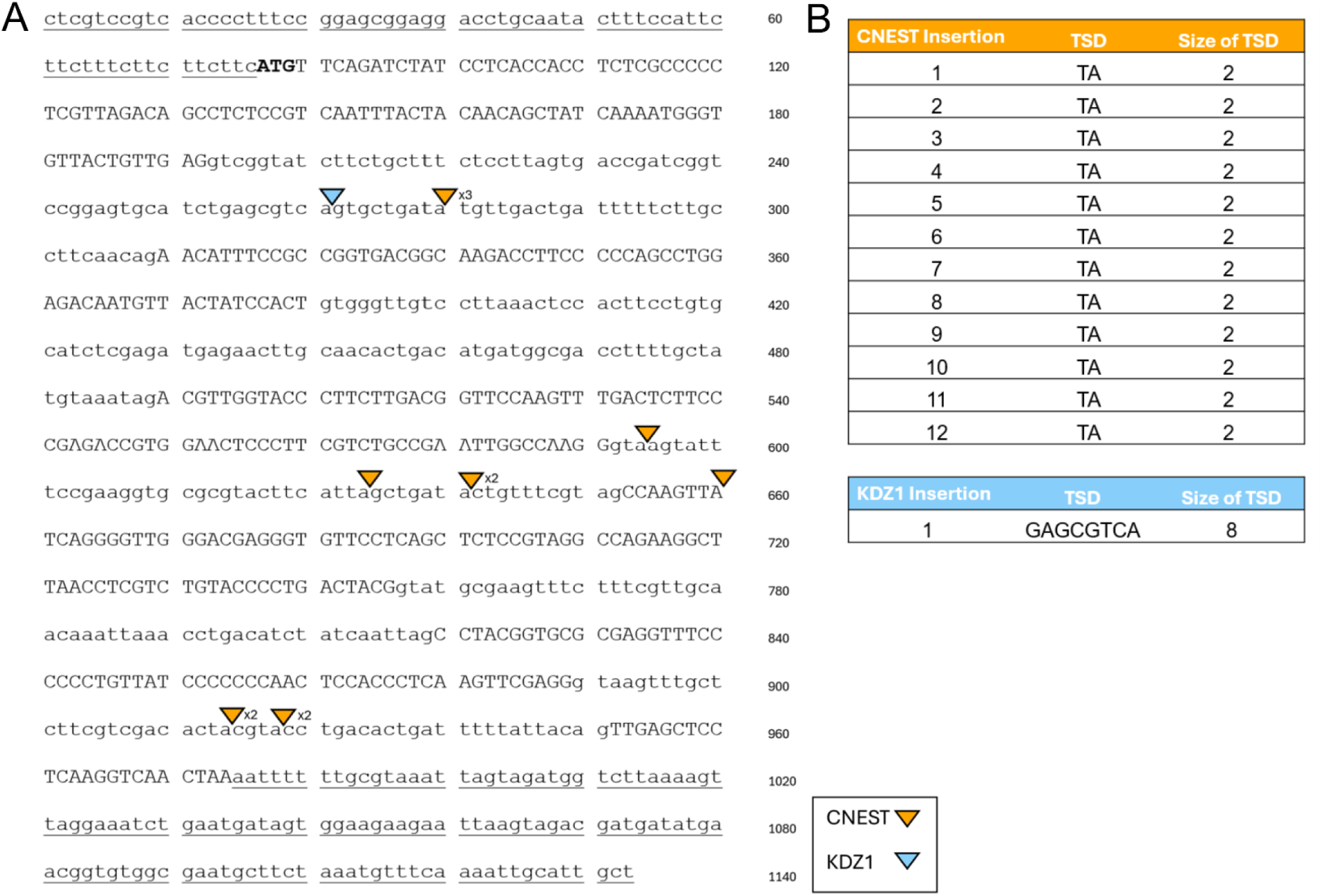
CNEST and KDZ1 insertion into *FRR1*. (**A**) A subset of CNEST (gold) and KDZ1 (light blue) insertion sites are mapped with arrows in *FRR1*. The 5′ and 3′ UTRs of *FRR1* are underlined; introns are shown in lowercase, exons in capital letters, and the start codon is bolded. (**B**) TSDs detected at CNEST and KDZ1 insertion sites. Insertions are ordered based on their appearance in the *FRR1* sequence from beginning to end.

**Fig S6.**
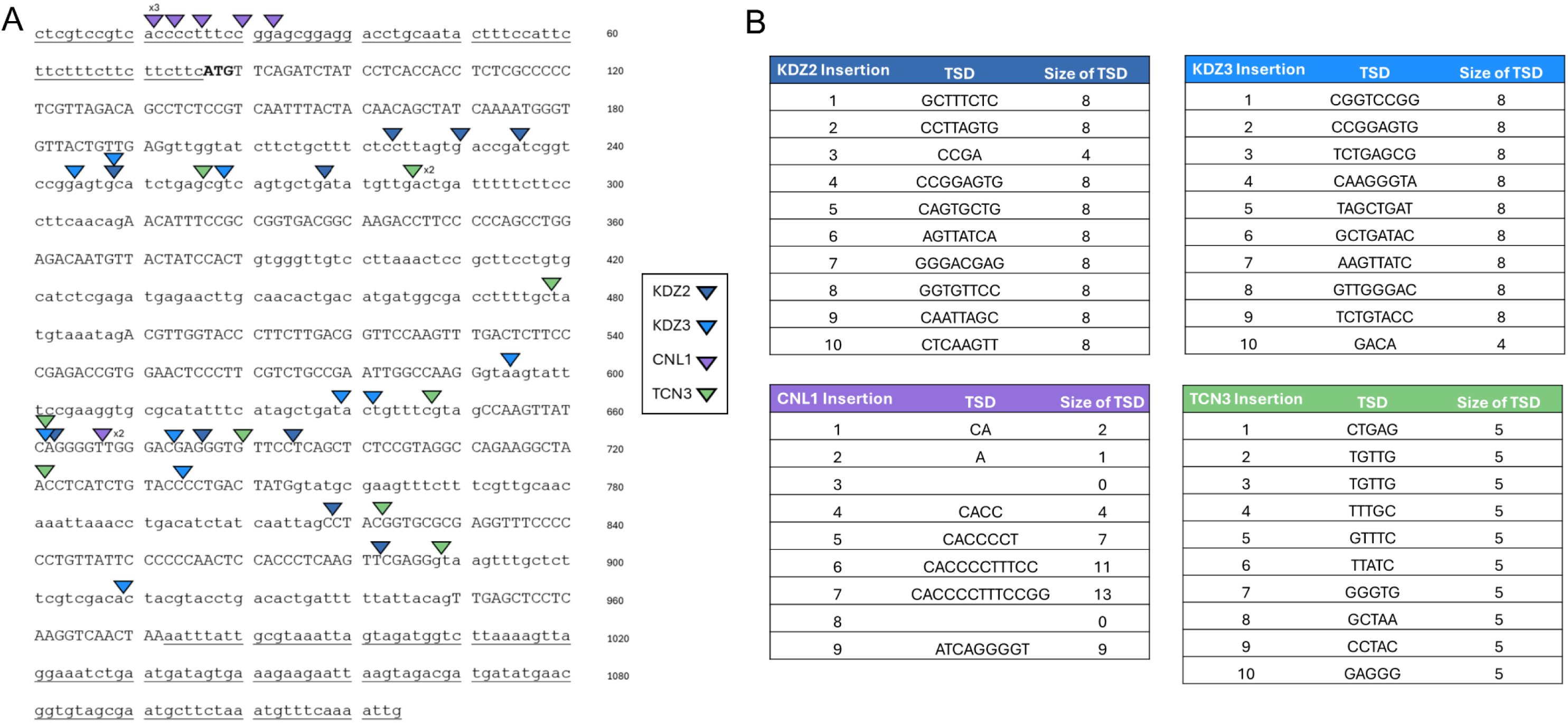
KDZ2, KDZ3, CNL1, and TCN3 insertion into *FRR1*. (**A**) A subset of KDZ2 (dark blue), KDZ3 (blue), CNL1 (purple), and TCN3 (green) insertion sites are mapped with arrows in *FRR1*. The 5′ and 3′ UTRs of *FRR1* are underlined; introns are shown in lowercase, exons in capital letters, and the start codon is bolded. (**B**) TSDs detected at KDZ2, KDZ3, CNL1, and TCN3 insertion sites. Insertions are ordered based on their appearance in the *FRR1* sequence from beginning to end.

**Fig S7.**
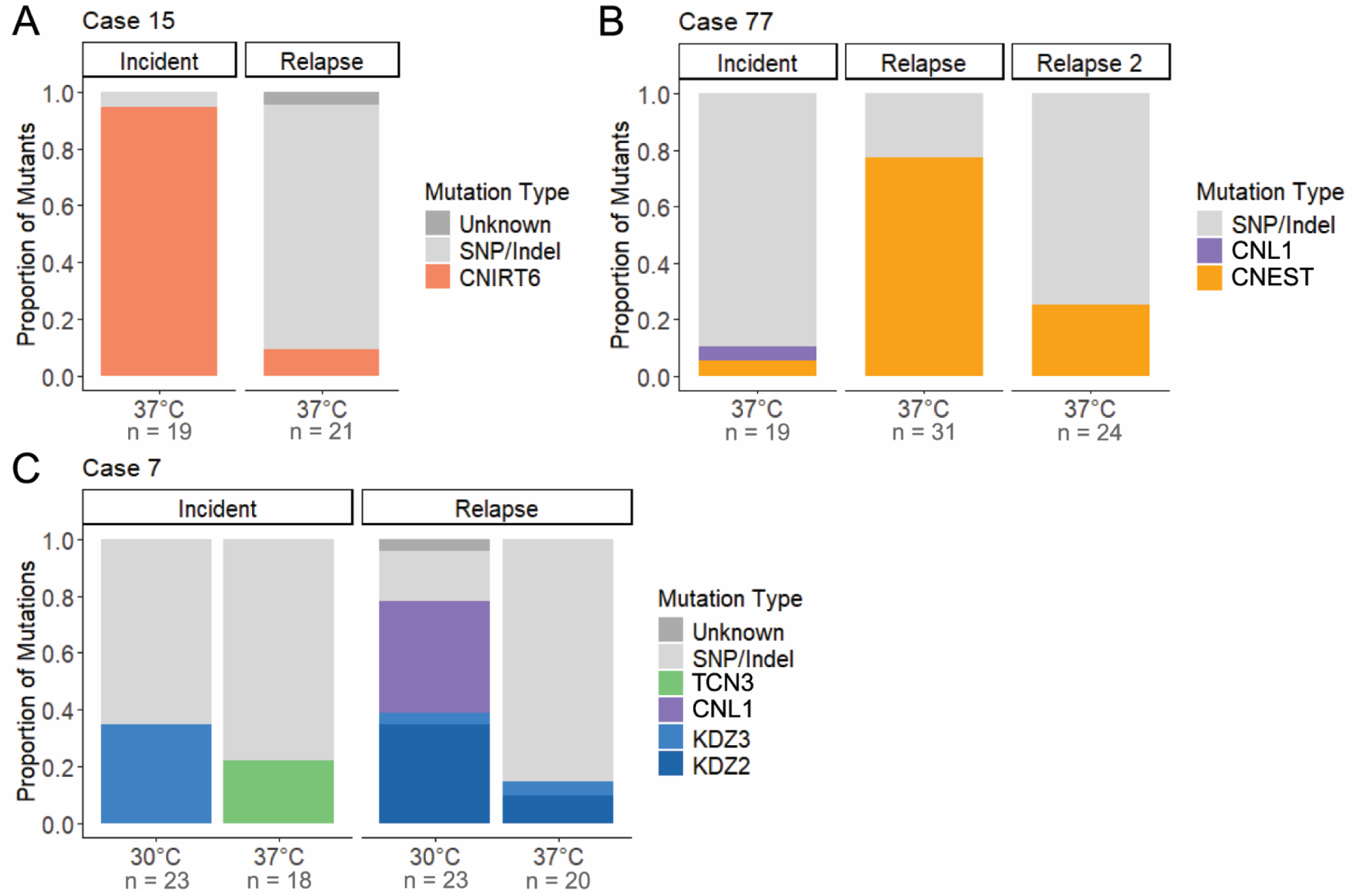
Growth in RPMI medium alters TE mobilization dynamics. Mutation spectra in the *FRR1* gene for independent rap+FK506-resistant mutants in case 15 (**A**), case 77 (**B**), and case 7 (**C**). The mutation type “Unknown” indicates failure to amplify *FRR1* or absence of detectable mutations in its sequence (N = 18 – 31).

**Fig S8.**
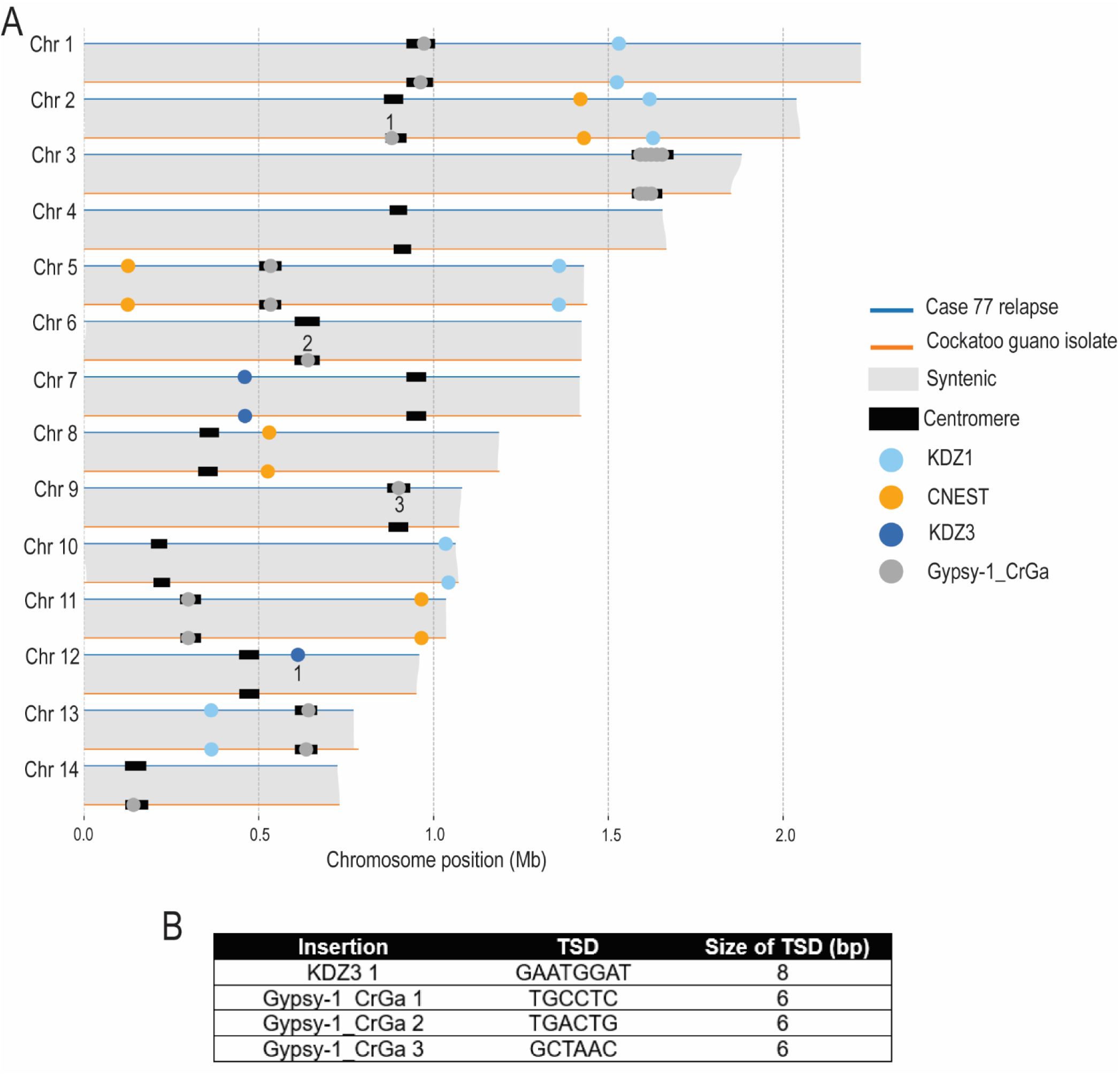
Transposition events between the cockatoo guano and case 77 relapse isolates. (**A**) Chromosome-level synteny between the case 77 relapse and cockatoo guano isolates. Full-length copies of CNEST (gold), KDZ1 (light blue), KDZ3 (dark blue), and Gypsy-1_CrGa (dark grey) are shown as circles. Predicted centromere regions are indicated by black rectangles and detected transposition events are numbered. (**B**) TSDs detected at KDZ3 and Gypsy-1_CrGa insertion sites.

**Fig S9.**
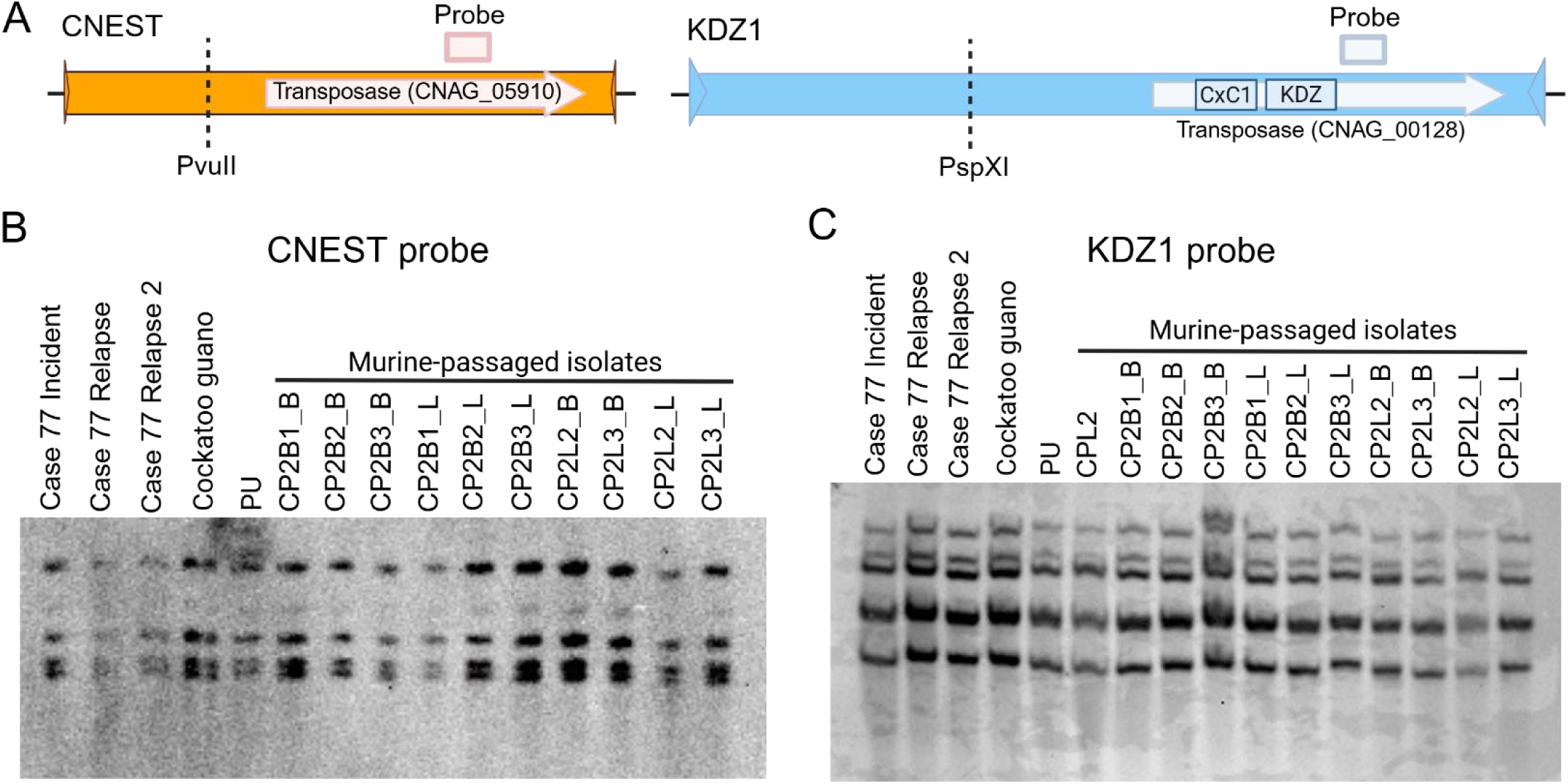
No predicted transposition of CNEST or KDZ1 in murine-passaged isolates. (**A**) Location of the restriction sites (dotted lines) and CNEST- and KDZ1-specific probes used in the Southern analysis. Southern blots of (**B**) PvuII-digested genomic DNA probed for CNEST, and (**C**) PspXI-digested genomic DNA probed for KDZ1. Shown are case 77 isolates, the cockatoo guano isolate, the patient isolate (PU) infected by the guano point-source strain, and cockatoo guano isolates passaged one to two times through mice by Sephton-Clark et al. (2023), recovered from brain or lung tissue (see Table S1 for strain details).

**Fig S10.**
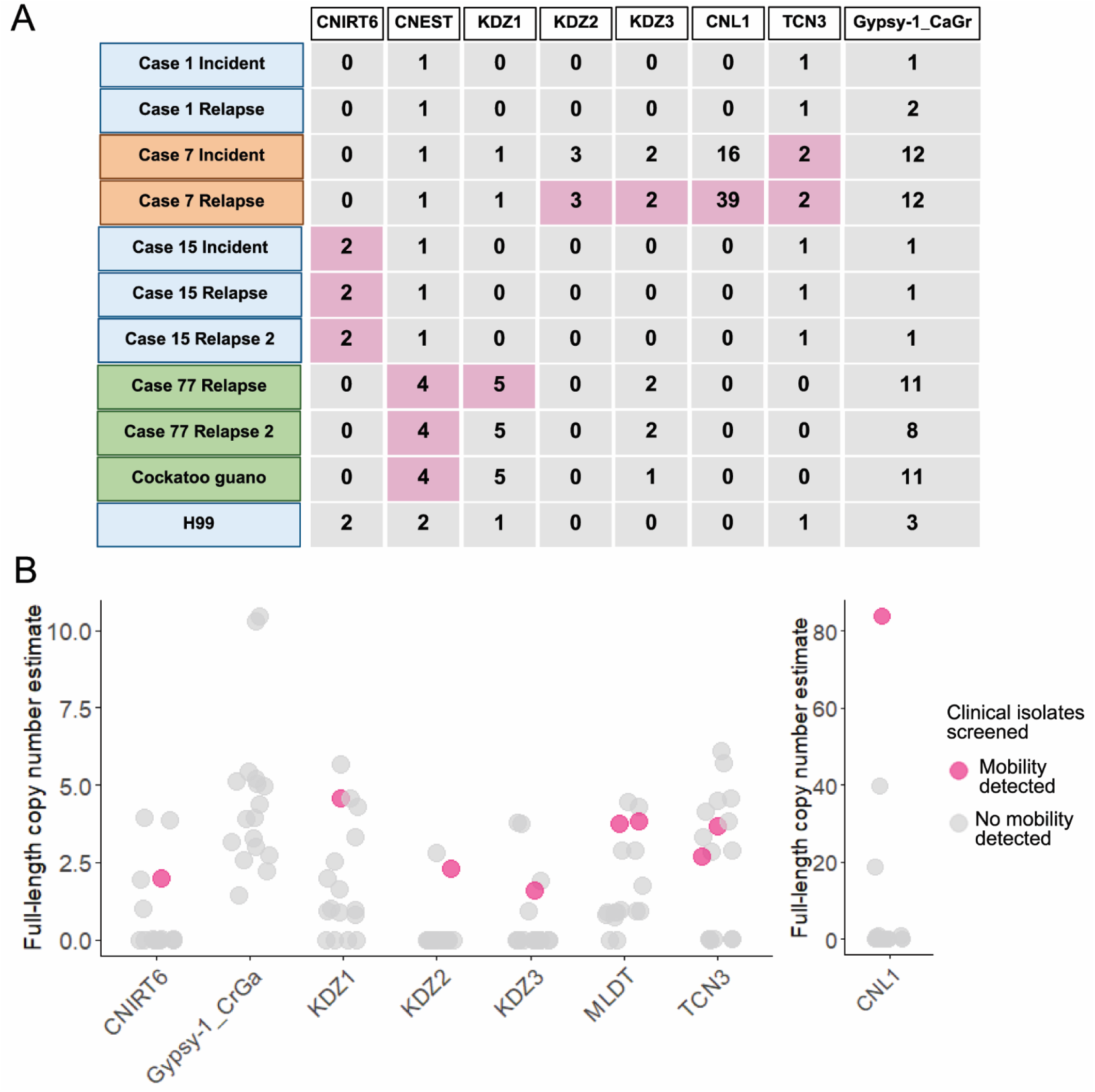
Full-length TE copies are present in genomes where no in vitro mobilization was detected. (**A**) Full-length TE copy numbers identified in whole-genome assemblies using BLAT and BLASTn homology searches. TE copy numbers in isolates with observed mobilization events into *FRR1* in YPAD medium are highlighted in pink. Isolate names are colored by lineage: VNI (blue), VNII (green), and VNBII (orange). (**B**) Full-length TE copy number estimates from short-read sequencing for all 15 *C. neoformans* incident isolates screened, except for case 7, where both incident and relapse isolates are shown. Isolates with no insertions into *FRR1* in YPAD medium are shown in grey, and those with insertions are shown in pink.

**Fig S11.**
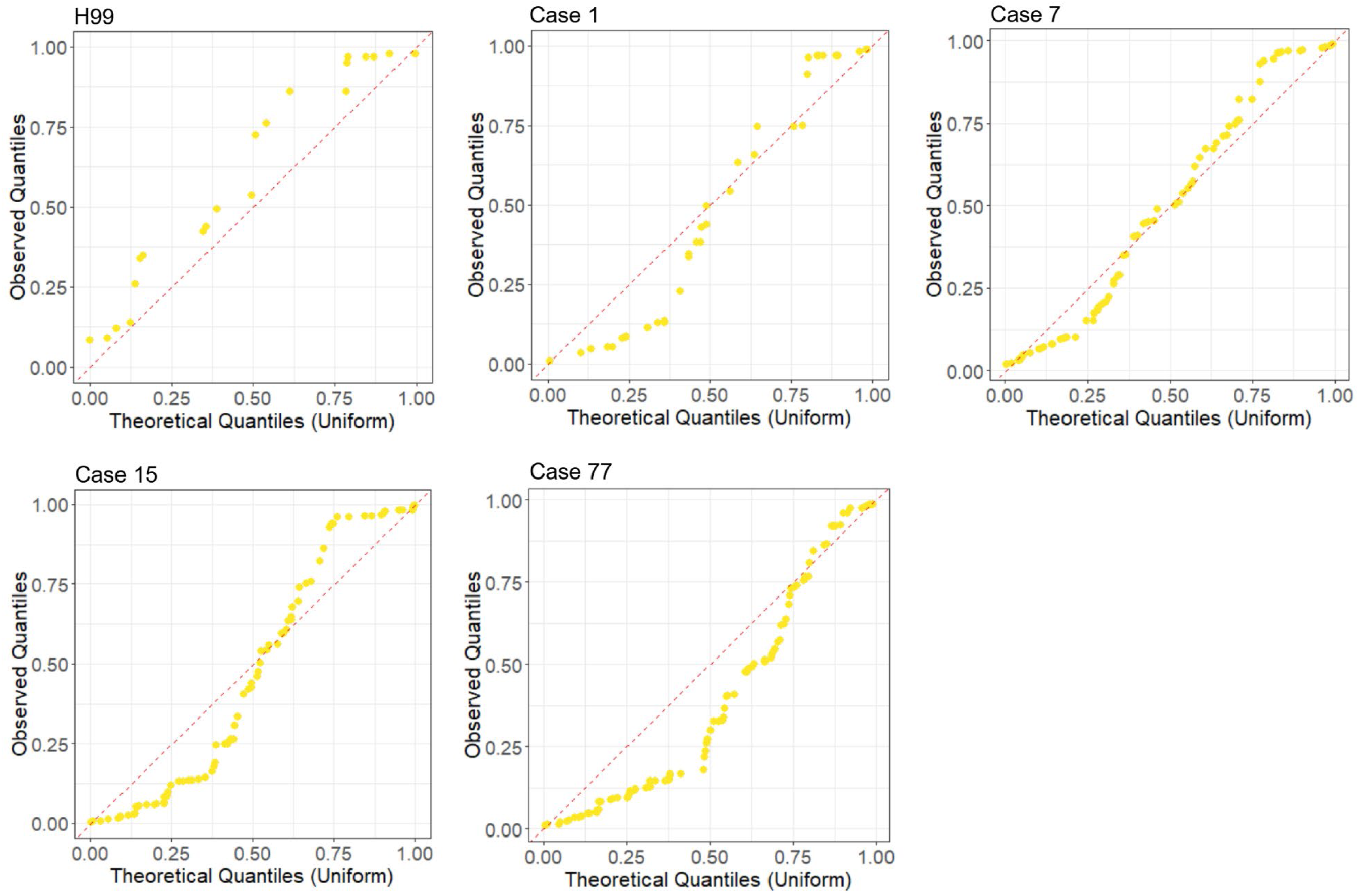
DNA TEs are approximately uniformly distributed across the genome. Quantile-quantile (QQ) plots are shown for the H99 reference genome and the first genome assembly from serially collected clinical isolates for cases 1, 7, 15, and 77 (representative examples). Observed quantiles correspond to the scaled positions (0 to 1) of DNA TEs along all chromosome arms. Theoretical quantiles are uniformly distributed random values between 0 and 1, sorted in ascending order, with their total number matching the number of observed DNA TEs.

**Fig S12.**
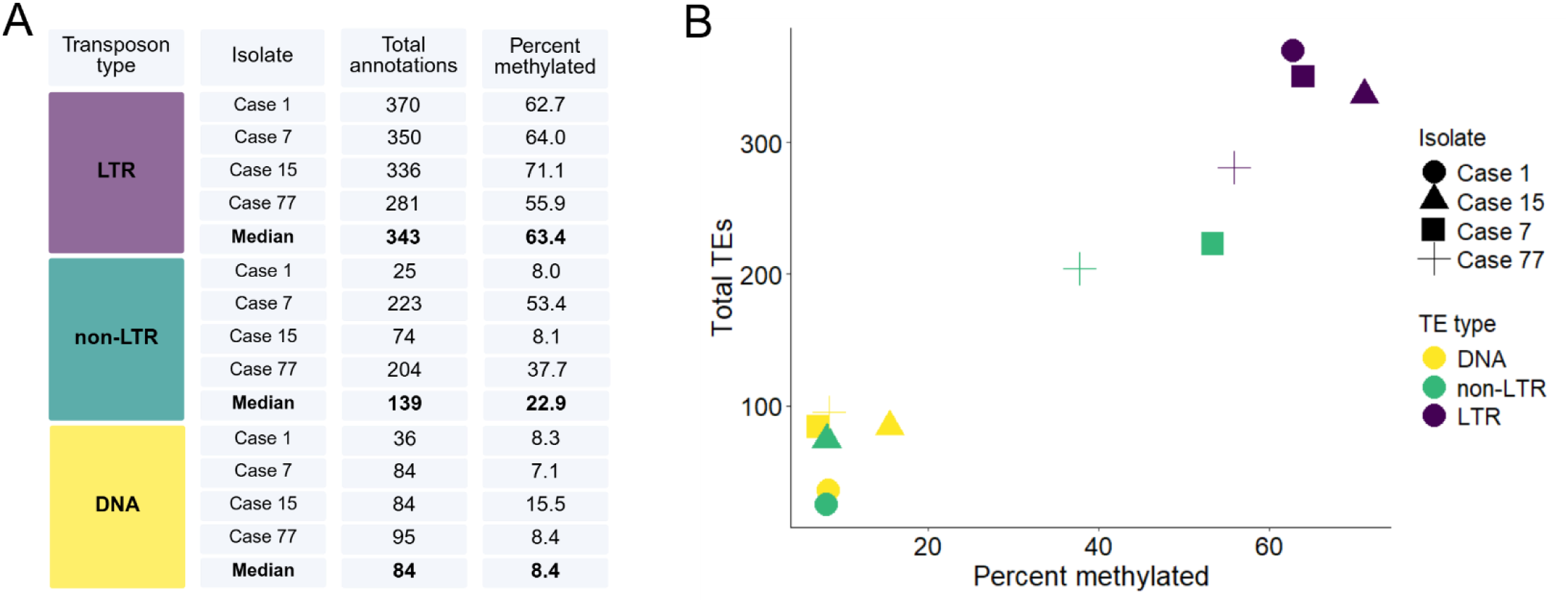
DNA transposons are less frequently methylated than LTR and non-LTR retrotransposons. (**A**) Number of TEs per type and the percentage containing at least one high-confidence DNA methylation call (methylation frequency > 0.75), with median values shown. (**B**) TE types (colors) per genome (shapes) and the percentage with at least one high-confidence DNA methylation call (methylation frequency > 0.75). The first genome assembly from serially collected clinical isolates was chosen as a representative.

**Table S1. Strains used in this study.**

**Table S2. Transposable elements characterized in this study.**

**Table S3. Whole-genome sequencing and assemblies used in this study.**

**Table S4. DNA methylation analysis of full-length transposable element copies.**

**Table S5. Primers used in this study.**

## Notes

### Competing Interest Statement

The authors have declared no competing interest.

### Summary of Updates

This revised version of the manuscript incorporates edits to the text and figures.

### Funding

- National Institutes of Health: R35GM118077, T32AI052080, R35GM122550
- National Institute of Allergy and Infectious Diseases: K99/R00AI166094, AI-73896, AI-93257, R01AI133654

